# BELLA: Berkeley Efficient Long-Read to Long-Read Aligner and Overlapper

**DOI:** 10.1101/464420

**Authors:** Giulia Guidi, Marquita Ellis, Daniel Rokhsar, Katherine Yelick, Aydın Buluç

**Author notes:** **Availability**: https://github.com/giuliaguidi/bella.

## Abstract

Recent advances in long-read sequencing enable the characterization of genome structure and its intra- and inter-species variation at a resolution that was previously impossible. Detecting overlaps between reads is integral to many long-read genomics pipelines, such as *de novo* genome assembly. While longer reads simplify genome assembly and improve the contiguity of the reconstruction, current long-read technologies come with high error rates. We present Berkeley Long-Read to Long-Read Aligner and Overlapper (BELLA), a novel algorithm for computing overlaps and alignments via sparse matrix-matrix multiplication that balances the goals of recall and precision, performing well on both.

We present a probabilistic model that demonstrates the feasibility of using short *k*-mers for detecting candidate overlaps. We then introduce a notion of *reliable k-mers* based on our probabilistic model. Combining *reliable k-mers* with our *binning* mechanism eliminates both the *k*-mer set explosion that would otherwise occur with highly erroneous reads and the spurious overlaps from *k*-mers originating in repetitive regions. Finally, we present a new method based on Chernoff bounds for separating true overlaps from false positives using a combination of alignment techniques and probabilistic modeling. Our methodologies aim at maximizing the balance between precision and recall. On both real and synthetic data, BELLA performs amongst the best in terms of F1 score, showing performance stability which is often missing for competitor software. BELLA’s F1 score is consistently within 1.7% of the top entry. Notably, we show improved *de novo* assembly results on synthetic data when coupling BELLA with the Miniasm assembler.

## 1 Introduction

Recent advancements in long-read sequencing technologies enable the characterization of genome structure and its variation between and within species that were not possible before. Nevertheless, the analysis of data after sequencing remains a challenging task. One of the biggest challenges for the analysis of high-throughput sequencing DNA fragments, namely *reads*, is whole genome assembly (Zhang *et al.*, 2011), which is the process of aligning and merging DNA fragments to reconstruct the original sequence. More specifically, *de novo* genome assembly reconstructs a genome from redundantly sampled reads without prior knowledge of the genome, enabling the study of previously uncharacterized genomes (Simpson and Durbin, 2012).

Long-read technologies (Eid *et al.*, 2009; Goodwin *et al.*, 2015) generate long reads with average lengths reaching and often exceeding 10,000 base pairs (bp). These allow the resolution of complex genomic repetitions, enabling more accurate ensemble views that were not possible with previous short-read technologies (Phillippy *et al.*, 2008; Nagarajan and Pop, 2009). However, the improved read length of these technologies comes at the cost of lower accuracy, with error rates ranging from 5% to 35%. Nevertheless, errors are more random and more evenly distributed within Pacific Biosciences long-read data (Giordano *et al.*, 2017) compared to short-read technologies.

The majority of state-of-the-art long-read assemblers uses the Overlap-Layout-Consensus (OLC) paradigm (Berlin *et al.*, 2015). The first step in OLC assembly consists of detecting overlaps between reads to construct an overlap (or string) graph. The OLC paradigm benefits from longer reads as significantly fewer reads are required to cover the genome, limiting the size of the overlap graph. Highly-accurate overlap detection is a major computational bottleneck in OLC assembly (Myers, 2014), mainly due to the compute-intensive nature of pairwise alignment.

At present, several algorithms are capable of overlapping error-prone long-read data with varying accuracy. The prevailing approach is to use an indexing data structure, such as a *k*-mer index table or a suffix array to identify a set of initial candidate read pairs, thus mitigating the high cost of computing pairwise alignments in a second stage (Chu *et al.*, 2016).

The process of identifying a set of initial candidate read pairs, sometimes simply known as *overlapping*, affects both the accuracy and the algorithm runtime. Precise identification of initial candidate read pairs minimizes the pairwise alignment running time while retaining all pairs that truly overlap in the genome. Computationally efficient and highly accurate overlapping and alignment algorithms have the potential to improve existing long-read assemblers, enabling *de novo* reference assemblies, detection of genetic variations of higher quality, and accurate metagenome classification. Our main contributions are:

1. Using a Markov chain model (Markov, 1971), we demonstrate the soundness of using a *k*-mer seed-based approach for accurately identifying initial candidate read pairs.
2. We develop a simple procedure for pruning *k*-mers and prove that it retains nearly all true overlaps with high probability. The result is greater computational efficiency without loss of accuracy.
3. We reformulate the problem of overlap detection in terms of a sparse matrix-matrix multiplication (SpGEMM), which enables the use of high-performance techniques not previously applied in the context of long read overlap and alignment.
4. Coupling our overlap detection with our newly developed seed-and-extend alignment algorithm, we introduce a novel method to separate true alignments from false positives.

## 2 Proposed Algorithm

The current work develops a computationally efficient and highly accurate algorithm for overlap detection and alignment for long-read genomics pipelines. The algorithm is implemented in a high-performance software package, called Berkeley Long-Read to Long-Read Aligner and Overlapper (BELLA).

BELLA uses a seed-based approach to detect overlaps in the context of long-read applications. Such an approach parses the reads into *k*-mers (i.e. sub-strings of fixed length *k*), which are then used as feature vectors to identify overlaps amongst all reads. Using a Markov chain model, we first show the feasibility of using a *k*-mer seed based approach for overlap detection of long-read data with high error rates.

Importantly, not all *k*-mers are created equal in terms of their usefulness for detecting overlaps. For instance, the overwhelming majority of *k*-mers that occur only once in the data set are errors (and are also not useful for seeding overlaps between pairs of reads). Similarly, *k*-mers that occur more frequently than what would be expected given the sequencing depth and the error rate are likely to come from repetitive regions. It is a common practice to prune the *k*-mer space using various methodologies (Koren *et al.*, 2017; Lin *et al.*, 2016; Carvalho *et al.*, 2016).

BELLA implements a novel method for filtering out *k*-mers that are likely to either contain errors or originate from a repetitive region. The *k*-mers that are retained by BELLA are considered to be *reliable*, where the reliability of a *k*-mer is defined as its probability of having originated from a unique (non-repetitive) region of the genome. BELLA’s reliable *k*-mer detection maximizes the retention of *k*-mers that belong to unique regions of the genome, using a probabilistic analysis given the error rate and the sequencing depth.

BELLA uses a sparse matrix to internally represent its data, where the rows are reads, columns are reliable *k*-mers, and a nonzero **A**(*i, j*) ≠ 0 contains the position of the *j*th reliable *k*-mer within *i*th read. Construction of this sparse matrix requires efficient *k*-mer counting.

Overlap detection is implemented in BELLA using SpGEMM, which allows our algorithm to achieve fast overlapping without using approximate approaches. SpGEMM is a highly flexible and efficient computational paradigm that enables the better organization of computation and generality because it can manipulate complex data structures such as the ones used in finding overlaps using shared *k*-mers. The implementation of this method within our pipeline enables the use of high-performance techniques previously not applied in the context of long-read alignment. It also allows continuing performance improvements in this step due to the ever-improving optimized implementations of SpGEMM (Nagasaka *et al.*, 2019; Deveci *et al.*, 2017).

BELLA’s overlap detection has been coupled with our high-performance seed-and-extend algorithm, meaning the alignment between two reads starts from a shared seed (identified in the previous overlap detection) and not necessarily from the beginning of reads. To refine the seed choice, we introduce a procedure, called *binning*. The *k*-mer positions in a read pair are used to estimate the length of the overlap, and the *k*-mers are “binned” based on their length estimates. We consider for alignment only *k*-mers belonging to the most crowded bins, termed *consensus k*-mers. During the alignment stage, BELLA uses a new method to separate true alignments from false positives depending on the alignment score. We prove that the probability of false positives decreases exponentially as the length of overlap between reads increases.

Existing tools also implement approximate overlap detection using sketching. A sketch is a reduced space representation of a sequence. Multiple randomized hash functions convert *k*-mers into integer fingerprints and a subset of these is selected to represent the sketch of a sequence according to some criterion. For example, Berlin *et al.* (2015) retain only the smallest integer for each hash function and use the collection of these minimum valued fingerprints as sketch. These methods, while fast, are approximate because sketching is a lossy transformation. Conversely, BELLA uses an explicit *k*-mer representation, which allows us to couple our overlap detection with a seed-and-extend alignment to refine the output and to improve the precision of our algorithm.

## 3 Methods

### Overlapping Feasibility

Chaisson and Tesler (2012) proposed a theory for how long reads contain subsequences that may be used to anchor alignments to the reference genome. The sequences are modeled as random processes that generate error-free regions whose length is geometrically distributed, with each such region separated by an error (Giordano *et al.*, 2017). The result obtained from their theory is the minimum sequence length to have an *anchor* within a confidence interval.

Here, we present an alternative model on how these subsequences, also known as *k*-mers, can be used to anchor alignments between two erroneous long reads, allowing an accurate overlap discovery among all the reads in a data set. The initial assumption of our model defines the probability of correctly sequencing a base as equal to *p* = (1 *− e*), where *e* is the error rate of the sequencer. From this notion, we model the probability of observing *k* correct consecutive bases on both *read*_1_ and *read*_2_ as a Markov chain process (Markov, 1971).

The Markov chain process is characterized by a *transition matrix* **P** that includes the probabilities to move from one state to another. Each row-index *start* of **P** represents the starting state, and each column-index *end* of **P** represents the ending state. Each entry of **P** is a non-negative number indicating a *transition probability*. Our transition matrix has (*k* + 1) possible states, which lead to (*k* + 1)^2^ transition probabilities of moving from *start* to *end*. The probability of having one correct base on both reads is *p*^2^. For any state except the *absorbing* state *k*, an error in at least one of the two sequences sets the model back to state 0, which happens with probability 1 − *p*^2^; otherwise, the Markov chain transition from state *i* to *i* + 1 happens with probability *p*^2^. The absorbing state *k* cannot be abandoned, as both *read*_1_ and *read*_2_ have already seen *k* consecutive correct bases. Hence, its transition probability is 1. Figure 1 describes the process: each state contains the number of successful sequenced bases obtained up to this point on both reads, while the arrows represent the transition probabilities.

**Figure 1:**
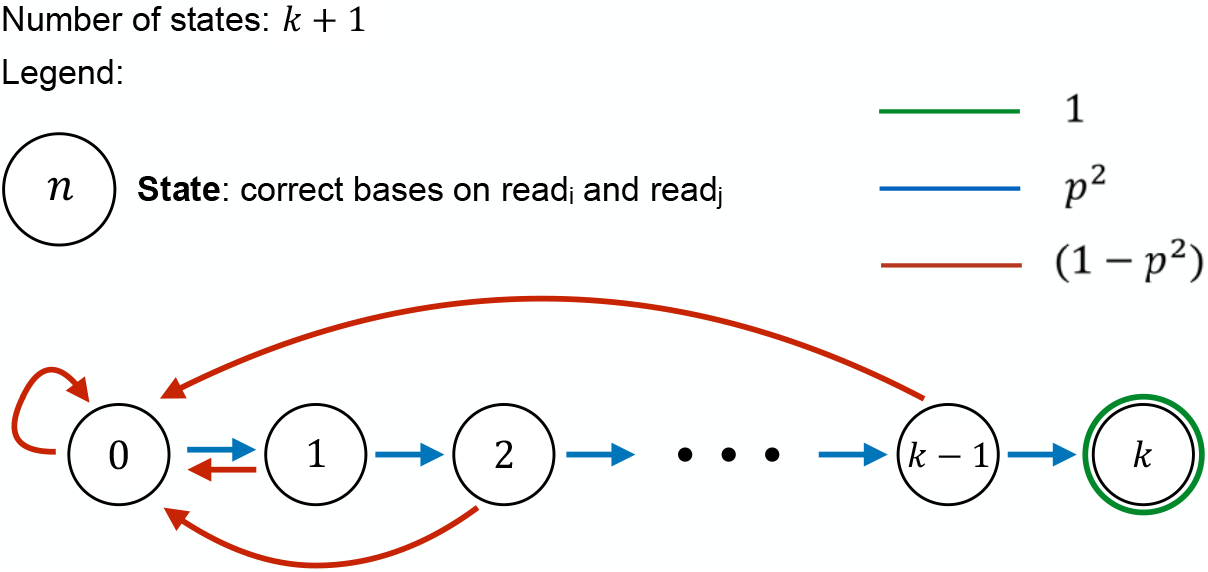
Proposed Markov chain model to prove the feasibility of usin short *k*-mers for overlap detection.

One can then find the probability of being in any of the states after *L* steps in the Markov chain by computing the *L*th power of the matrix **P**, where *L* is the length of the overlap between the two sequences. More efficiently, one can compute this iteratively using just *L* sparse matrix-vector products starting from the unit vector **v***←* (1, 0, …, 0), as shown in Algorithm 1. This approach is sufficient because we are only interested in the probability of being in the final absorbing state. The above operation leads to the probability of having one correct *k*-mer in the same location on both reads given a certain overlap region.

#### Algorithm 1

Probability of observing at least one shared correct *k*-mer in an overlap region of length L *> k*.

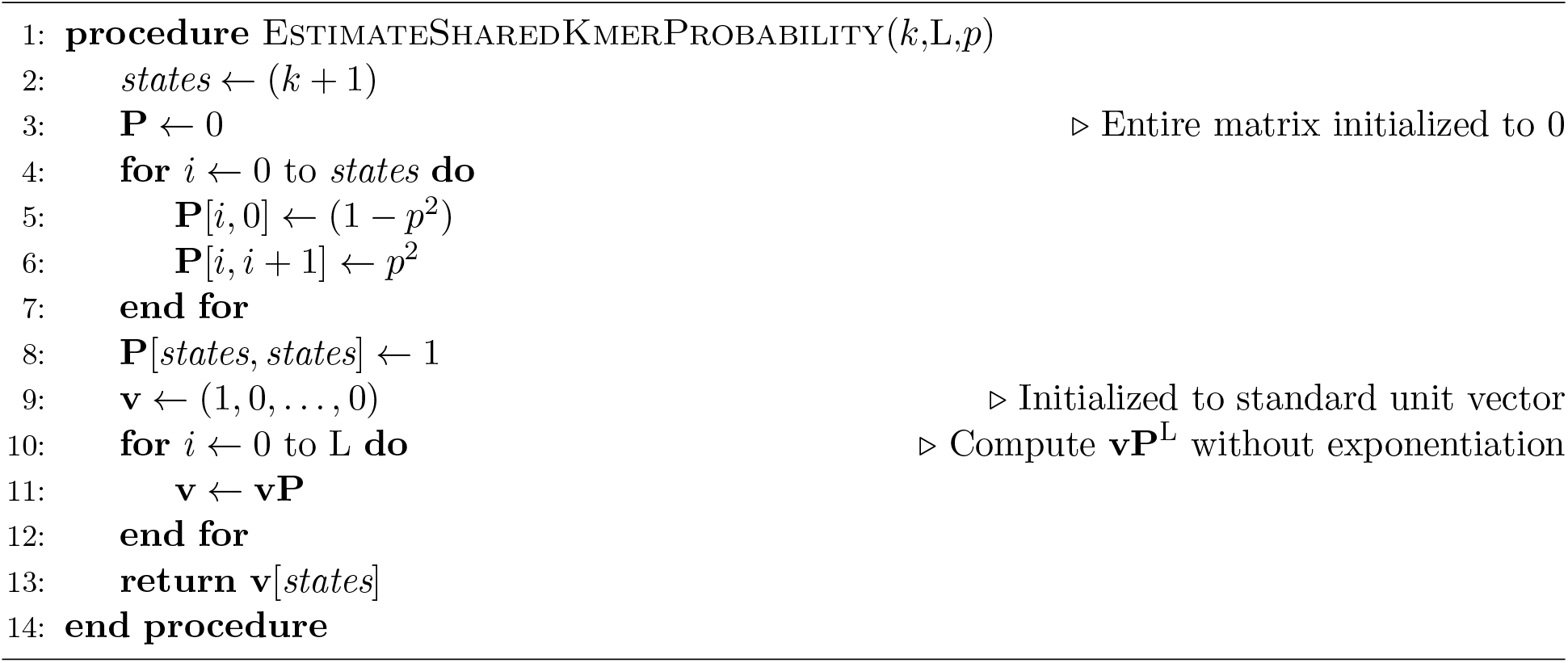

The proposed model is the driving factor behind the choice of the optimal *k*-mer length to be used during overlap detection.

### Reliable *k*-mers

Repetitive regions of the genome lead to certain *k*-mers occurring frequently in the input reads. *k*-mers from these regions pose two problems for pairwise overlapping and alignment. First, their presence increases the computational cost, both at the overlapping stage and at the alignment stage, because these *k*-mers generate numerous and possibly spurious overlaps. Second, they often do not provide valuable information.

Our argument here is that *k*-mers coming from a repetitive region in the genome can be ignored for seed-based overlapping. This is because either (a) the read is longer than the repeat, in which case there should be enough sequence data from the non-repeat section to find overlaps, or (b) the read is shorter than the repeat, in which case their overlaps are ambiguous and uninformative to begin with and will not be particularly useful for downstream tasks such as *de novo* assembly.

Following the terminology proposed by Lin *et al.* (2016), we identify *k*-mers that do not exist in the genome as *non-genomic*, thus characterizing *k*-mers present in the genome as *genomic*. A genomic *k*-mer can be *repeated*, if it is present multiple times in the genome, or *unique*, if it is not. One can think of the presence of *k*-mers within each read as that read’s feature vector. For the reasons discussed above, the feature vector should include all the unique *k*-mers, as they often are the most informative features.

Since we do not know the genome before assembly, we must estimate the genomic uniqueness of *k*-mers from our redundant, error-containing reads. In this section, we provide a mathematically grounded procedure that chooses a frequency range for *k*-mers that we consider being *reliable*. The basic question that guides the reliable *k*-mer selection procedure is the following: “Given that a *k*-mer is sequenced from a unique (non-repeat) region of the genome, what is the probability it will occur at least *m* times in the input data?”. For a genome *G* sequenced at depth *d*, the conditional modeled probability is:

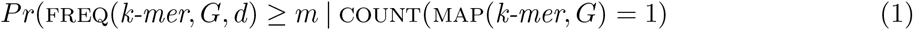

where MAP(*k-mer, G*) is the set of locations in the genome *G* where *k-mer* can be mapped, COUNT() function computes the cardinality of a given input set, and FREQ(*k-mer, G, d*) is the expected number of occurrences of *k-mer* within sequenced reads, assuming each region of *G* is sequenced *d* times (*sequencing depth*). In that sense, BELLA’s approach to select reliable *k*-mers diverges sharply from how Lin *et al.* (2016) selects their *solid strings*. While solid strings discard infrequent *k*-mers, our model discards highly-repetitive *k*-mers, arguing that (a) unique *k*-mers are sufficient to find informative overlaps, and (b) a unique *k*-mer has a low probability of occurring frequently.

The probability of a *k*-mer being sequenced correctly is approximately (1 *− e*)^*k*^, where *e* is the error rate. The probability of correctly sequencing a *k*-mer once can be generalized to obtain the probability of seeing it multiple times in the data, considering that each correct sequencing of that *k*-mer is an independent event. For example, if the sequencing depth is *d*, the probability of observing a unique *k*-mer *k*_*i*_ in the input data *d* times is approximatively (1 *− e*)^*dk*^. More generally, the number of times a unique *k* length section of the genome is sequenced correctly when the sequencing depth is *d* follows a binomial distribution:

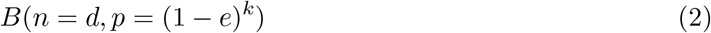

where *n* is the number of trials and *p* is the probability of success. Consequently, we derive that the probability of observing a *k*-mer *k*_*i*_ (which corresponds to a unique, non-repetitive region of the genome) *m* times within a sequencing input data with depth *d* is:

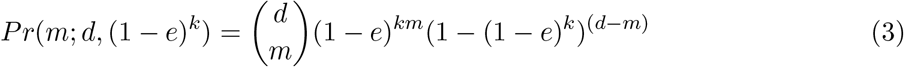

where *m* is the multiplicity of the *k*-mer in the input data, *e* is the error rate, *d* is the sequencer depth, and *k* is the *k*-mer length. Given the values of *d*, *e*, and *k*, the curve *Pr*(*m*; *d,* (1 *− e*)^*k*^) can be computed.

Equation 3 is used to identify the range of reliable *k*-mers. To select the lower bound *l*, we compute *Pr*(*m*; *d,* (1 *− e*)^*k*^) for each multiplicity *m* and cumulatively sum up these probabilities, starting from *m* = 2. The cumulative sum does not start from *m* = 1 because a *k*-mer appearing a single time in the input data (and therefore appearing on a single read) cannot be used to identify the overlap between two reads. The lower bound *l* is the smaller *m* value after which the cumulative sum exceeds a user-defined threshold *ϵ*. The choice of *l* matters when the sequencing error rate is relatively low (*≈* 5%) or when the sequencing coverage is high (*≈* 50 *−* 60*×*), or both. This is because in those cases, a *k*-mer with small multiplicity has a high probability to be incorrect.

The upper bound *u* is chosen similarly. Here, the probabilities are cumulatively summed up starting from the largest possible value of *m* (i.e. *d*). In this case, *u* is the largest value of *m* after which the cumulative sum exceeds the user-defined threshold *ϵ*. The *k*-mers that appear more frequently than *u* have too low a probability of belonging to a unique region of the genome, and multi-mapped *k*-mers would lead to an increase of computational cost, and potentially to misassemblies.

*K*-mers appearing with greater multiplicities than *u* and those appearing with smaller multiplicities than *l* in the input set are discarded and not used as read features in the downstream algorithm. Our reliable *k*-mer selection procedure discards at most 2*ϵ* useful information in terms of *k*-mers that can be used for overlap discovery.

### Sparse Matrix Construction and Multiplication

BELLA uses a sparse matrix format to store its data and sparse matrix-matrix multiplication (SpGEMM) to identify overlaps. Sparse matrices express the data access patterns concisely and clearly, allowing better organization of computation. The sparse matrix **A**, also known as the *data matrix*, is a *|reads|*-by-*|k-mers|* matrix with reads as its rows and the entries of the *k*-mer dictionary as its columns. If the *j*th reliable *k*-mer is present in the *i*th read, the cell (*i, j*) of **A** is non-zero. **A** is then multiplied by its transpose, **A**^T^, yielding a sparse *overlap matrix* **AA**^T^ of dimensions *|reads|*-by-*|reads|*. Each non-zero cell (*i, j*) of the overlap matrix contains the number of shared *k*-mers between the *i*th read and the *j*th read and the corresponding positions in the corresponding read pair of (at most) two shared *k*-mers.

The column-by-column sparse matrix multiplication is implemented efficiently using the Compressed Sparse Columns (CSC) format for storing sparse matrices. However, other options are certainly possible in the future, which is one of the advantages of our work. Any novel sparse matrix format and multiplication algorithm would apply to the overlapping problem and would enable continued performance improvements since multiple software packages already implement this primitive, including Intel MKL and Sandia’s KokkosKernels (Deveci *et al.*, 2017).

The SpGEMM algorithm shown in Figure 2 is functionally equivalent to a *k*-mer based seed-index table, which is common in other long-read alignment codes. However, the CSC format allows true constant-time random access to columns as opposed to hash tables. More importantly, the computational problem of accumulating the contributions from multiple shared *k*-mers to each pair of reads is handled automatically by the choice of appropriate data structures within SpGEMM. Figure 2 illustrates the merging operation of BELLA, which uses a hash table data structure indexed by the row indexes of **A**, following the multi-threaded implementation proposed by Nagasaka *et al.* (2019). Finally, the contents of the hash table are stored into a column of the final matrix once all required nonzeros for that column are accumulated.

**Figure 2:**
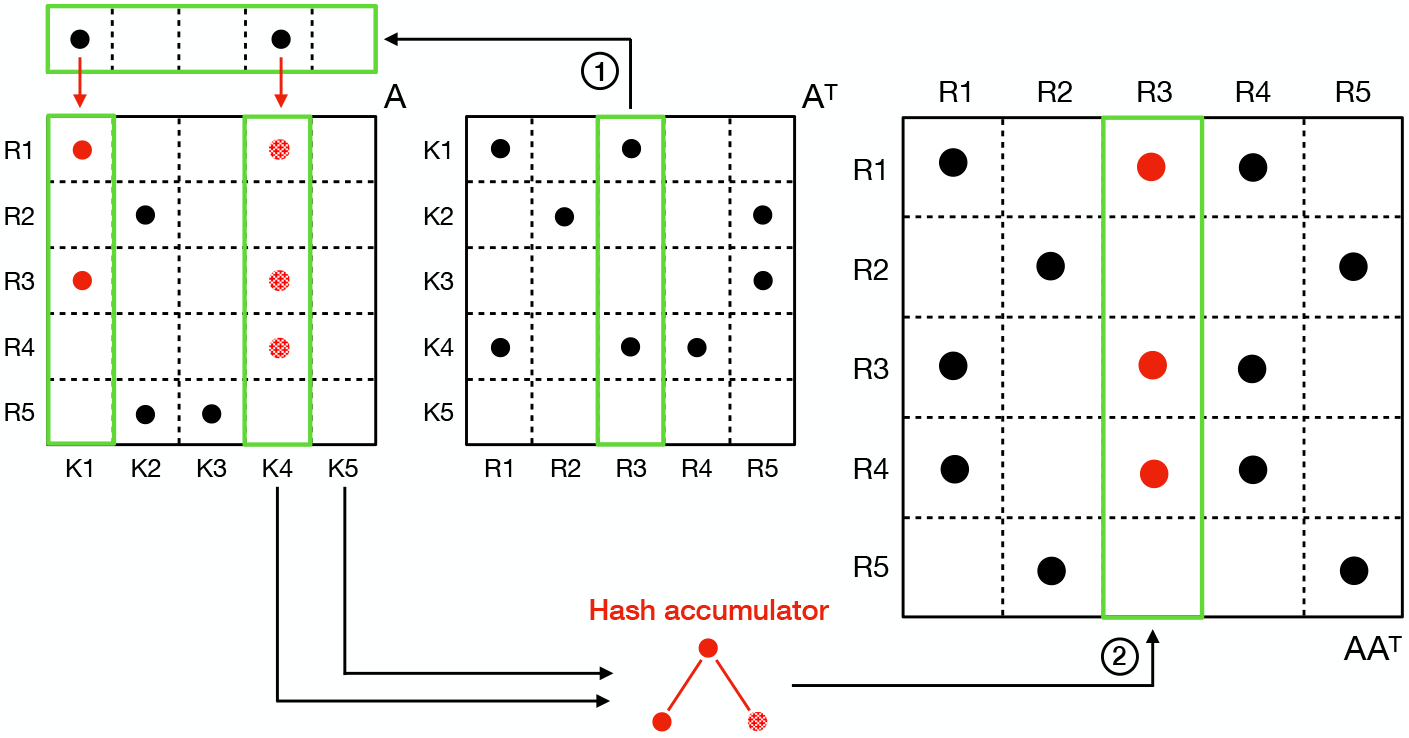
Column-by-column sparse matrix multiplication. **A**^T^(:*, R*_3_) is selected as the “active column”: its non-zero elements define which columns of **A** need to be considered, by looking at their corresponding row indexes in **A**^T^.

Since BELLA performs a seed-and-extend alignment from pairs of reads that share at least *t* (by default *t* = 1) *k*-mers, the overlapping stage needs to keep track of the positions of the shared *k*-mers. In the cases of multiple *k*-mers shared between any pair of reads, it is not economical to store all *k*-mer matches. Furthermore, while the reliable *k*-mer selection procedure described earlier eliminates most of the hits from repetitive regions, it is still possible to observe misleading *k*-mer matches due to sequencing errors and repetitions.

To ensure an optimal seed choice for the alignment step, BELLA employs the following binning methodology. Whenever a shared *k*-mer is encountered between a pair of reads, we use its location in these reads to estimate an overlap length and orientation. A new overlap estimate forms its bin with a single element unless it is already within the boundaries of a previous bin (i.e. within an adjustable distance *β*, which is 500 by default), in which case that nearby bin’s vote count is incremented by one as long as the two originating *k*-mers are not overlapping. Only two bins with the highest number of votes are retained, together with a representative *k*-mer supporting that overlap estimate.

In the resulting sparse overlap matrix **AA**^T^, each non-zero cell (*i, j*) is a structure composed of an integer value storing the number of shared *k*-mers, and an integer array of size 4 storing the position on read *i* and on read *j* of (up to two) shared *k*-mers corresponding to the overlap estimate bins with highest number of votes. To enable this special multiplication that performs scalar multiplication and additions differently, we use the semiring abstraction (Kepner and Gilbert, 2011). Multiplication on a semiring allows the user to overload scalar multiplication and addition operations and still use the same SpGEMM algorithm. Many existing SpGEMM implementations support user-defined semirings, including those that implement the GraphBLAS API (Buluç *et al.*, 2017).

Increasing the genome size also increases the memory requirements for building the final overlap matrix. For large genomes, it is possible that the sparse overlap matrix **AA**^T^ would not fit in memory even if the data matrix **A** does. BELLA avoids this situation by dividing the multiplication into batches based on the available RAM. At each stage, only a batch of columns of the overlap matrix are created. The set of nonzeros in that batch of the overlap matrix are immediately tested for alignments (as described in Section 3). The pairs that pass the alignment test are written to the output file of BELLA so that the current batch of overlap matrix can be discarded.

Given the nature of our problem, the sparse overlap matrix **AA**^T^ is a symmetric matrix. Thus, we compute the multiplication using only the lower triangle of **A**, avoiding computing the pairwise alignment twice for each pair. Currently, there are no known specialized SpGEMM implementations for **AA**^T^ that store and operate only on **A**, but we hope to develop one in the future. This would have cut the memory requirements in half. The obvious solution of computing inner projects of rows of **A** is suboptimal, because it has to perform Ω(*|reads|*^2^) inner products even though the majority of inner products are zero. By contrast, our column-by-column implementation runs faster than *O*(*|reads|*^2^) whenever the overlap matrix **AA**^T^ is sparse. Given that the main purpose of the overlapping process is used to filter candidate pairs, the overlap matrix tends to be very (over 99%) sparse in practice.

### Pairwise Alignment

As high precision is desirable for avoiding wasted work in subsequent stages of *de novo* assembly, BELLA filters candidate read pairs by performing fast, near linear-time pairwise seed-and-extend alignments.

As opposed to approaches that rely on sketches or minimizers, such as Minimap2 and MHAP, seed-and-extend alignment can be performed directly using the *k*-mers from BELLA’s overlap stage. BELLA’s alignment module is based on our high-performance seed-and-extend banded-alignment, which uses a narrow adaptive band that appreciably improves performance reducing the search space for the optimal alignment.

Our binning mechanism returns at most two seed *k*-mers, which are used as inputs to the seed-and-extend x-drop alignment. For each read pair in the overlap matrix, the alignment is extended from one- or two-seed *k*-mers until the alignment score drops *x* points below the best score seen so far. Once the alignment is complete, if the best score is lower than a threshold *n*, the pair of sequences is discarded.

Since a fixed alignment score threshold might not capture true alignments, we use an *adaptive threshold*, calculated according to the estimated overlap between a given pair of reads. The choice of the scoring matrix used in the pairwise alignment step can justify the alignment score threshold being a linear function of the estimated overlap length.

Given an estimated overlap region of length *L* and the probability *p* = *q*^2^ of getting a correct base on both sequences, we would expect *m* = *p · L* correct matches within that overlap region. The alignment score *χ* can be written as follows:

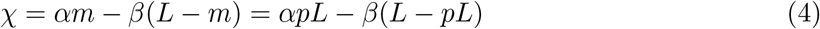

where *m* is the number of matches, *L* is the length of the overlap region, *α* is the value associated with a match in the scoring matrix while *β* is the penalty for mismatch or a gap/indel (*α, β >* 0). Given these assumptions, we define the ratio *φ* between *χ* and the estimated overlap length *L* as:

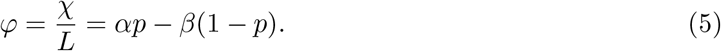

The expected value of *φ* is equal to 2 *· p −* 1, if an exact alignment algorithm is used. We would like to define a cutoff in the form of (1 *− δ*)*φ*, so that we retain pairs over this cutoff as true alignments and discard remaining pairs. We use a Chernoff bound (Chernoff *et al.*, 1952; Hoeffding, 1963) to define the value of *δ*, proving that there is only a small probability of missing a true overlap of length *L ≥* 2000 bp (which is the minimum overlap length for a sequence to be considered a true-positive) when using the above-defined cutoff. We derived the following Chernoff bound:

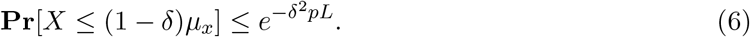

Given two sequences that indeed overlap by *L* = 2000, the probability of their alignment score being below the mean by more than 10% (*δ* = 0.1) is *≤* 5.30 *×* 10^*−*7^. The derivation of the above formula is reported in Supplementary Material. BELLA achieved high values of recall and precision among state-of-the-art software tools, with an x-drop value of *x* = 50 and an adaptive threshold derived from the scoring matrix and the with *δ* = 0.1 cutoff rate.

## 4 Evaluation

Table 1 lists the data sets used for evaluation. The selected genomes have varying size and complexity since analysis results are sensitive to these features (Li *et al.*, 2012).

**Table 1:** Data sets used for evaluation. Data sets above the line are real data, while data sets below the line have been generated using PBSIM (Ono *et al.*, 2012). Download: portal.nersc.gov/project/m1982/bella/

| Short Name | Species and Strain | Fastq Size |
| --- | --- | --- |
| *E.coli* (Sample) 30X | *Escherichia coli* MG1655 strain | 266 MB |
| *E.coli* 100X | *Escherichia coli* MG1655 strain | 929 MB |
| *C.elegans* 40X | *Caenorhabditis elegans* Bristol mutant strain | 8.90 GB |
| *P. aeruginosa* 30X | *Pseudomonas aeruginosa* PAO1 | 359 MB |
| *V. vulnificus* 30X | *Vibrio vulnificus* YJ016 | 288 MB |
| *A. baumannii* 30X | *Acinetobacter baumannii* | 248 MB |
| *C.elegans* 20X | *Caenorhabditis elegans* | 3.75 GB |

As for performance metrics, we use recall, precision, F1 score, and running time. The recall is defined as the fraction of true positives of the aligner/overlapper over the total size of the *ground truth*; precision is the fraction of true positives of the aligner/overlapper over the total number of elements found by the aligner/overlapper. F1 score is the harmonic average of precision and recall.

We consider a read pair as true-positive if the sequences align for at least 2 kb in the reference genome. We derived the threshold *t* = 2 kb from the procedure proposed by Li (2016), and generated the ground truth using Minimap2. A detailed description of our evaluation procedure and ground truth generation can be found in Supplementary Material.

We also report preliminary assembly results on simulated data sets, obtained by coupling the overlappers/aligners with the Miniasm assembler (Li, 2016). As assembly quality metrics, we include the number of contigs, the number of misassemblies, N50, and the total assembly length. A contig is defined as a set of overlapping reads that together represent a consensus region of the genome. Misassemblies are the set of positions in the contigs whose flanking sequences map more than 1kbps away from each other. N50 is a measure of the assembly contiguity and is defined as the minimum contig length needed to cover 50% of the genome.

## 5 Results

BELLA is evaluated against several state-of-the-art software for long-read overlap detection and alignment, using both synthetic and real PacBio data sets. The synthetic data sets were generated using PBSIM (Ono *et al.*, 2012) with an error rate of 15%. Notably, the advantage of synthetic data is that the ground truth is known. Table 2 and Table 3 show results on synthetic and real data sets, respectively, in terms of accuracy and runtime. The last column of each table indicates whether the respective overlapper also performs nucleotide-level alignment on overlapping reads. Table 4 illustrates assembly results on simulate data sets. Each overlapper was coupled with Miniasm assembler (Li, 2016).

**Table 2:** Recall, precision, F1 score, and time comparison (synthetic data). The last column indicates if the tool computes alignments. Bold font indicates best performance and underlined font indicates the second-best. DALIGNER did not run with synthetic data sets.

| Data Set | Overlapper | Recall | Precision | F1 Score | Time (s) | Alignment |
| --- | --- | --- | --- | --- | --- | --- |
| *P. aeruginosa* 30X | BELLA | <u>97.66</u> | 89.68 | <u>93.50</u> | 140.43 | Y |
|  | BLASR | 86.86 | <u>90.54</u> | 88.66 | 230.97 | Y |
|  | MECAT | 38.40 | **95.20** | 54.72 | <u>21.76</u> | N |
|  | Minimap2 | **99.10** | 88.83 | **93.69** | **17.76** | N |
|  | MHAP | 72.68 | 63.42 | 67.74 | 72.72 | N |
| *V. vulnificus* 30X | BELLA | **97.27** | 84.05 | <u>90.18</u> | 42.76 | Y |
|  | BLASR | 87.31 | 84.74 | 86.01 | 179.51 | Y |
|  | MECAT | 43.53 | <u>88.89</u> | 58.44 | <u>17.20</u> | N |
|  | Minimap2 | <u>96.71</u> | **89.33** | **92.87** | **12.93** | N |
|  | MHAP | 74.52 | 45.10 | 56.19 | 70.21 | N |
| *A. baumannii* 30X | BELLA | **97.54** | 84.90 | <u>90.78</u> | 44.69 | Y |
|  | BLASR | 89.51 | 84.58 | 86.98 | 152.76 | Y |
|  | MECAT | 46.31 | **90.39** | 61.25 | <u>15.65</u> | N |
|  | Minimap2 | <u>96.89</u> | <u>85.79</u> | **91.01** | **10.06** | N |
|  | MHAP | 76.88 | 28.79 | 41.89 | 69.02 | N |
| *C. elegans* 20X | BELLA | 91.80 | <u>88.02</u> | **89.87** | 2,352.24 | Y |
|  | BLASR | <u>95.61</u> | 78.19 | 86.02 | 3,655.12 | Y |
|  | MECAT | 13.45 | **95.09** | 23.57 | <u>222.10</u> | N |
|  | Minimap2 | **95.76** | 82.84 | <u>88.83</u> | **194.77** | N |
|  | MHAP | 82.57 | 6.41 | 11.90 | 5,353.30 | N |

**Table 3:** Recall, precision, F1 score, and time comparison (real data). The last column indicates whether the considered aligner does actual alignment or just overlap detection. BLASR result for *C. elegans* 40X is not reported as BLASR v5.1 does not accept `fastq` larger than 4 GB.

| Data Set | Overlapper | Recall | Precision | F1 Score | Time (s) | Alignment |
| --- | --- | --- | --- | --- | --- | --- |
| *E. coli* (Sample) | BELLA | 82.66 | **85.69** | **84.15** | 60.94 | Y |
|  | DALIGNER | <u>89.97</u> | 62.62 | 73.84 | **8.70** | Y |
|  | BLASR | 77.64 | <u>79.64</u> | 78.63 | 160.05 | Y |
|  | MECAT | 78.41 | 78.71 | 78.56 | 24.45 | N |
|  | Minimap2 | **91.40** | 76.36 | <u>83.21</u> | <u>16.57</u> | N |
|  | MHAP | 79.71 | 66.93 | 72.76 | 43.67 | N |
| *E. coli* | BELLA | 65.08 | 71.22 | <u>68.01</u> | 374.37 | Y |
|  | DALIGNER | <u>82.18</u> | 54.50 | 65.54 | **58.50** | Y |
|  | BLASR | 35.41 | <u>72.01</u> | 47.48 | 715.19 | Y |
|  | MECAT | 54.61 | **72.69** | 62.37 | <u>84.21</u> | N |
|  | Minimap2 | **80.68** | 62.30 | **70.30** | 107.76 | N |
|  | MHAP | 67.84 | 44.60 | 53.81 | 287.66 | N |
| *C. elegans* 40X | BELLA | 75.43 | <u>73.81</u> | **74.61** | 9,042.36 | Y |
|  | DALIGNER | 62.81 | 58.66 | 60.67 | 22,797.50 | Y |
|  | MECAT | 73.05 | **75.27** | <u>74.14</u> | **733.79** | N |
|  | Minimap2 | **94.13** | 34.06 | 50.02 | <u>1,733.26</u> | N |
|  | MHAP | 86.63 | 5.31 | 10.01 | <u>9,102.23</u> | N |

**Table 4:**
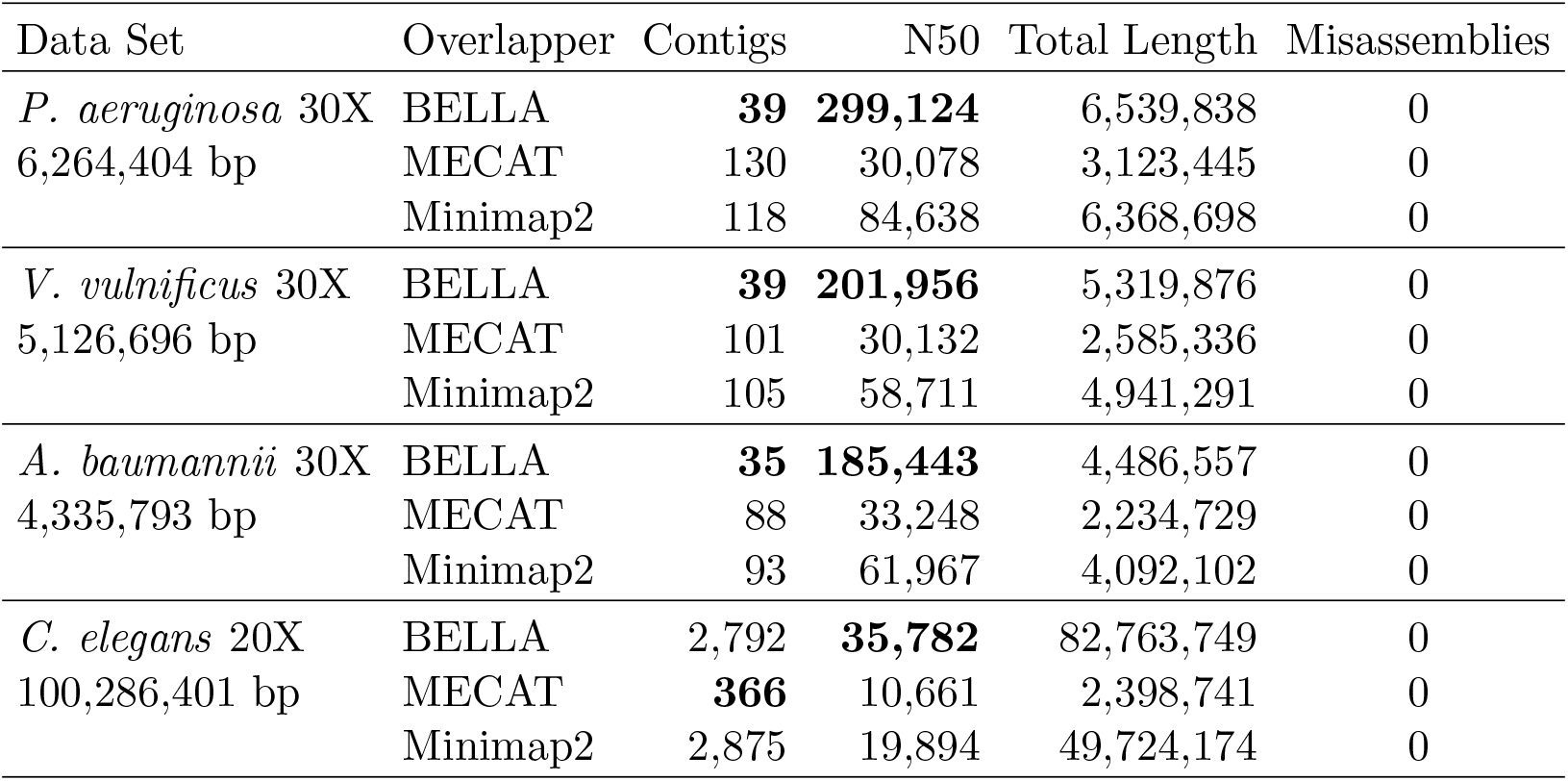
Preliminary assembly result of synthetic data sets. Overlappers’ outputs have been translated in PAF format and paired with Miniasm (Li, 2016) assembler. DALIGNER did not run with synthetic data sets. Miniasm did not produce any output when paired with BLASR and MHAP.

| Data Set | Overlapper | Contigs | N50 | Total Length | Misassemblies |
| --- | --- | --- | --- | --- | --- |
| *P. aeruginosa* 30X<br>6,264,404 bp | BELLA | **39** | **299,124** | 6,539,838 | 0 |
|  | MECAT | 130 | 30,078 | 3,123,445 | 0 |
|  | Minimap2 | 118 | 84,638 | 6,368,698 | 0 |
| *V. vulnificus* 30X<br>5,126,696 bp | BELLA | **39** | **201,956** | 5,319,876 | 0 |
|  | MECAT | 101 | 30,132 | 2,585,336 | 0 |
|  | Minimap2 | 105 | 58,711 | 4,941,291 | 0 |
| *A. baumannii* 30X<br>4,335,793 bp | BELLA | **35** | **185,443** | 4,486,557 | 0 |
|  | MECAT | 88 | 33,248 | 2,234,729 | 0 |
|  | Minimap2 | 93 | 61,967 | 4,092,102 | 0 |
| *C. elegans* 20X<br>100,286,401 bp | BELLA | 2,792 | **35,782** | 82,763,749 | 0 |
|  | MECAT | **366** | 10,661 | 2,398,741 | 0 |
|  | Minimap2 | 2,875 | 19,894 | 49,724,174 | 0 |

Table 2 shows that MECAT trades off recall for precision, achieving the highest precision but missing a large number of the true overlaps. In contrast, BELLA, Minimap2, and BLASR were consistently strong (typically over 80%) in both precision and recall, but BLASR had a much higher computational cost (2.6*×* slower than BELLA on average). BELLA’s F1 score is consistently higher than competitor software with the exception of Minimap2, which had a slight improvement of 1.1% on three out of four data sets, while BELLA had an improvement of 1.2% over Miniamp2 on *C. elegans* 20X. Minimap2 was the fastest tool for synthetic data, performing only overlapping and not alignment.

Table 3 shows that, although BLASR performed reasonably well on the synthetic data sets, it achieved lower recall than other software on real data sets. BLASR did not run on *C. elegans* 40X as its latest version (v5.1) does not accept `fastq` larger than 4 GB^1^. DALIGNER proved to be the fastest of the tools on *E. coli* 30X and *E. coli* 100X, but its performance decreases drastically when moving to a larger data set where DALIGNER has the worst running time, 2.5*×* slower than BELLA that also performs alignment. BELLA’s F1 score outperformed competitor software with the exception of Minimap2 on *E. coli* 100X.

Table 4 shows that BELLA significantly improved assembly quality with respect to competitor software. MECAT produced fewer contigs than BELLA on *C. elegans* 20X, but its N50 and assembly length are significantly smaller, meaning MECAT did not retain sufficiently many true overlaps to perform the assembly. Miniasm did not produce any assembly when coupled with MHAP and BLASR.

## 6 Discussion

BELLA proposes a computationally efficient and highly accurate approach for overlapping and aligning noisy long reads, based on mathematical models that minimize the cost of overlap detection while maximizing the retention of true overlaps. Tables 2 and 3 show BELLA’s competitive accuracy compared to state-of-the-art software, demonstrating the effectiveness of the methodologies we introduced and implemented within BELLA. Our runtime is within the average of competitive software, which is noteworthy given that BELLA performs nucleotide-level alignments that are sufficiently accurate to facilitate downstream analysis.

On synthetic data, BELLA achieves both high recall and precision, consistently among the best. On real data, recall and precision are generally lower than for synthetic data, nevertheless, BELLA’s F1 scores remain amongst the best, showing performance stability which is often missing in competitor software. Notably, BELLA has a 49.16% higher F1 score than Minimap2 for *C. elegans* 40X. Overall, a good performer on one data set becomes one of the worst on some other data set whereas BELLA’s F1 score is consistently within 1.7% of the top entry.

Tables 2 and 3 show that BELLA achieves higher values of F1 score on synthetic data compared to real data. The way ground truth is generated could explain such behavior. For synthetic data, the ground truth comes directly with the data set itself. Thus, we know the exact location from which a read originates in the reference genome and which other reads overlap with it. For real data, the read locations in the reference are determined by mapping the reads to the reference using Minimap2 in its “mapping mode”, as detailed in our Supplementary Material. Intuitively, such procedure is suboptimal as there is no guarantee that Minimap2 correctly locates every single read. BELLA could potentially find a better set of true overlaps than those identified by Minimap2. Given a uniformly covered genome, we observed that Minimap2 and other long read mappers tend to map reads to “hotspots” within a genome instead of mapping them uniformly across the genome. This results in uneven coverage and overestimation of overlaps by a factor of 1.14*×*, as shown in the Supplementary Material. Recall above a certain point on real data would mean the overlapper is overestimating the overlap cardinality as well. Hence, it is possible that BELLA’s true accuracy on real data is higher in reality. We plan to investigate these issues deeper in the future.

Table 4 gives an insight into the beneficial impact of BELLA as part of a *de novo* assembly pipeline. Our methodologies, such as the reliable *k*-mer procedure, sensibly improve the assembly contiguity when compared to MECAT and Minimap2. Importantly, Miniasm did not produce any output when using either MHAP or BLASR, which leads us to emphasize that an assembler is often built with a particular overlapping tool in mind, is programmed to take advantage of that tool’s methodologies. We plan to build our assembler upon BELLA to fully exploit its potential.

## 7 Conclusion

Long-read sequencing technologies enable highly accurate reconstruction of complex genomes. Read overlapping is a major computational bottleneck in long-read genomic analysis pipelines such as genome and metagenome assembly.

We presented BELLA, a computationally efficient and highly accurate long-read to long-read aligner and overlapper. BELLA uses a *k*-mer based approach to detect overlaps between noisy, long reads. We demonstrated the feasibility of the *k*-mer based approach through a mathematical model based on Markov chains. BELLA provides a novel algorithm for pruning *k*-mers that are unlikely to be useful in overlap detection and whose presence would only incur unnecessary computational costs. Our reliable *k*-mers detection algorithm explicitly maximizes the probability of retaining *k*-mers that belong to unique regions of the genome.

BELLA achieves fast overlapping without sketching using sparse matrix-matrix multiplication (SpGEMM), implemented utilizing high-performance software and libraries developed for this sparse matrix subroutine. Any novel sparse matrix format and multiplication algorithm would be applicable to overlap detection and enable continued performance improvements. We coupled BELLA’s overlap detection with our newly developed seed-and-extend banded-alignment algorithm. The choice of the optimal *k*-mer seed occurs through our binning mechanism, where *k*-mer positions within a read pair are used to estimate the length of the overlap and to “bin” *k*-mers to form a consensus.

We developed and implemented a new method to separate true alignments from false positives depending on the alignment score. This method demonstrates that the probability of false positives decreases exponentially as the overlap length between sequences increases.

BELLA achieves consistently high values of accuracy compared to state-of-the-art tools on both synthetic and real data, while being performance competitive. BELLA appreciably improves assembly results on synthetic data, validating our approach. Future work includes a further characterization of real data features, performance improvements, and the development of an assembler built on BELLA.

## Acknowledgements

We thank Heng Li for his help with Minimap2 and Miniasm, Rob Egan and Steven Hofmeyr for valuable discussions, and NECST Laboratory, Elizabeth Koning, and Ed Younis for key collaborations.

## Supplementary Material

### S1 Overlap Length Definition

Given two sequences *s*_1_ and *s*_2_ sharing a *k*-mer *k*_1,2_ of length *k* as illustrated in Figure 3, we first check if 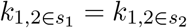 since BELLA stores only the lexicographical smaller *k*-mer between itself and its reverse complement. If 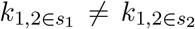, we reverse complement *s*_1_ and update 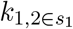 location accordingly,

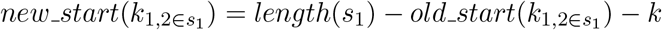

**Figure 3:**
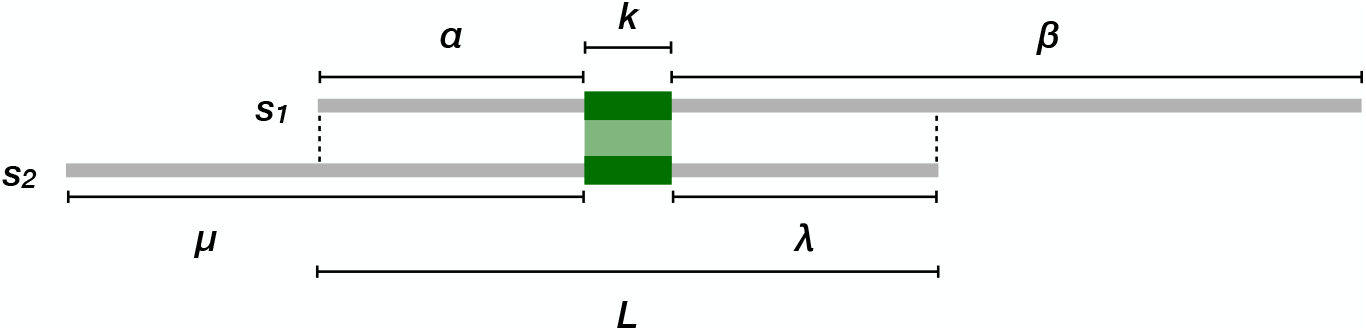
The overlap length estimate *L* is computed starting from a seed location as the sum of the seed length *k*, the smallest left margin (*α*), and the smallest right margin (*λ*).

Then, we define *α* as the starting position of *k*_1,2_ on *s*_1_, and, similarly, *μ* as the starting position of *k*_1,2_ on *s*_2_, *β* as the difference between the ending position of *k*_1,2_ on *s*_1_ and the length of *s*_1_, and *λ* as the difference between the ending position of *k*_1,2_ on *s*_2_ and the length of *s*_2_.

The overlap length estimate *L* is computed as follows,

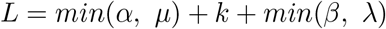

Notably, PacBio long-read data has high insertion rates. Nevertheless, errors are approximately evenly distributed within reads (Giordano *et al.*, 2017). Hence, we would expect roughly the same number of insertions in *s*_1_ and *s*_2_, within their overlap region. This implies our overlap approximation takes into consideration insertions and accounts for them, on average.

### S2 Overlapping Feasibility

Figure 4 illustrates the probabilities of finding one correct shared *k*-mer between two reads by varying the value of *k*, the error rate, and the overlap length *L*. The probability of a *k*-mer being correct decreases approximately geometrically as its length increases. With decreasing error rate, however, a larger *k* would be preferable since it would decrease the amount of *k*-mers coming from repetitive regions of the genome. For a given a minimum overlap length *τ*, our model favors the selection of the largest *k*-mer length with sufficiently high probability (*≈* 80%) to find a correct seed for an overlap of length *τ*, while not increasing the runtime by choosing an excessively small Notice that an overwhelming majority of the overlaps will be longer than *τ*, so the ultimate recall will be significantly higher than 80%.

**Figure 4:**
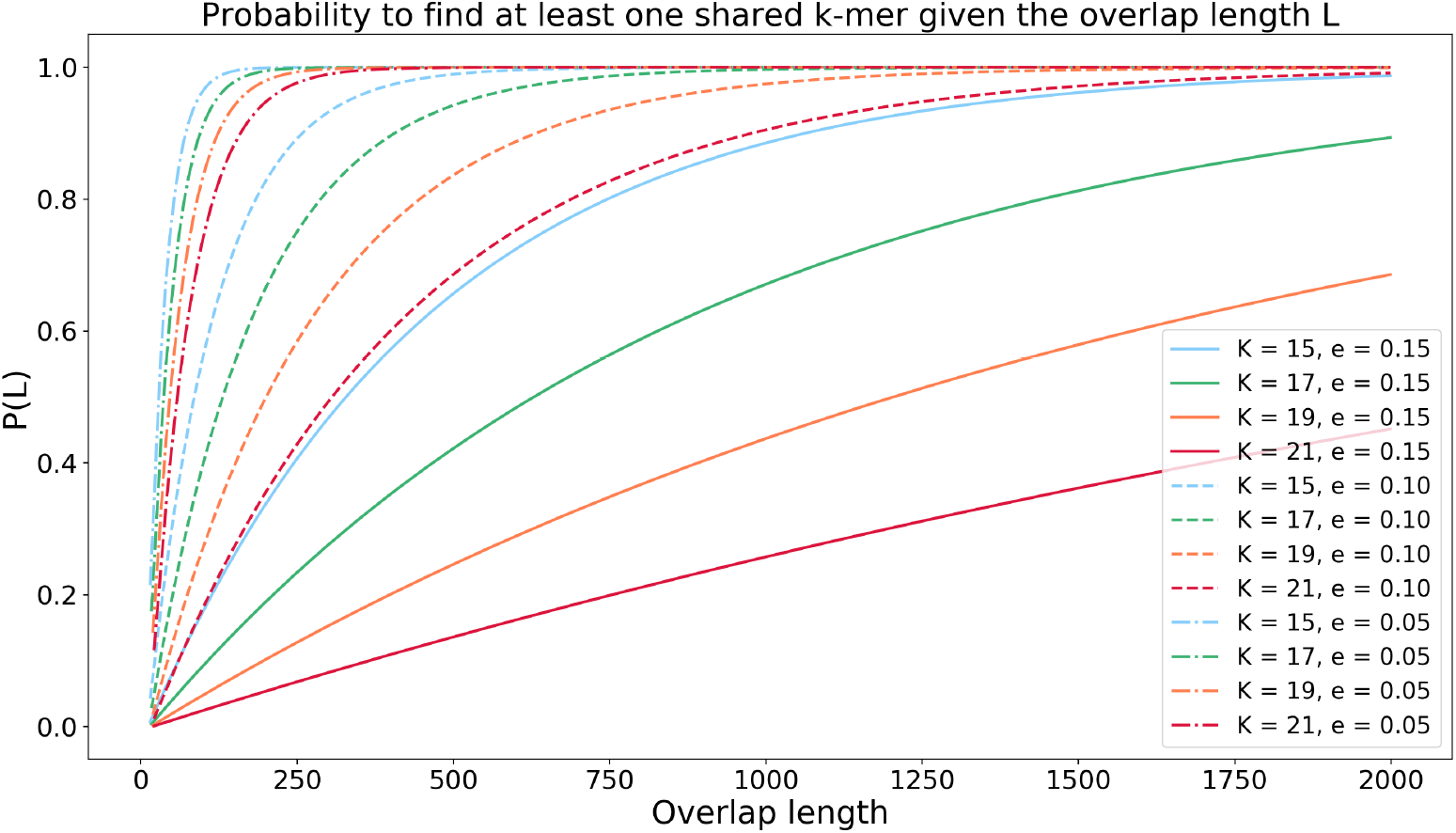
Outcome of the proposed model. The probability of success is set to *p* = 0.85, *p* = 0.90, and *p* = 0.95 representing error rates *e* of 0.15, 0.10, and 0.05.

Assuming that we are interested in an overlap of length *L* ≥ 2000 as defined in our ground truth^2^, our model in Figure 4 would suggest a *k*-mer length of 17 for a data set with error rate *≈* 15%. Figure 5 suggests that *k* = 17 offers a desirable trade-off between high recall and precision and low runtime for a data set generated with an error rate of *≈* 15%.

**Figure 5:**
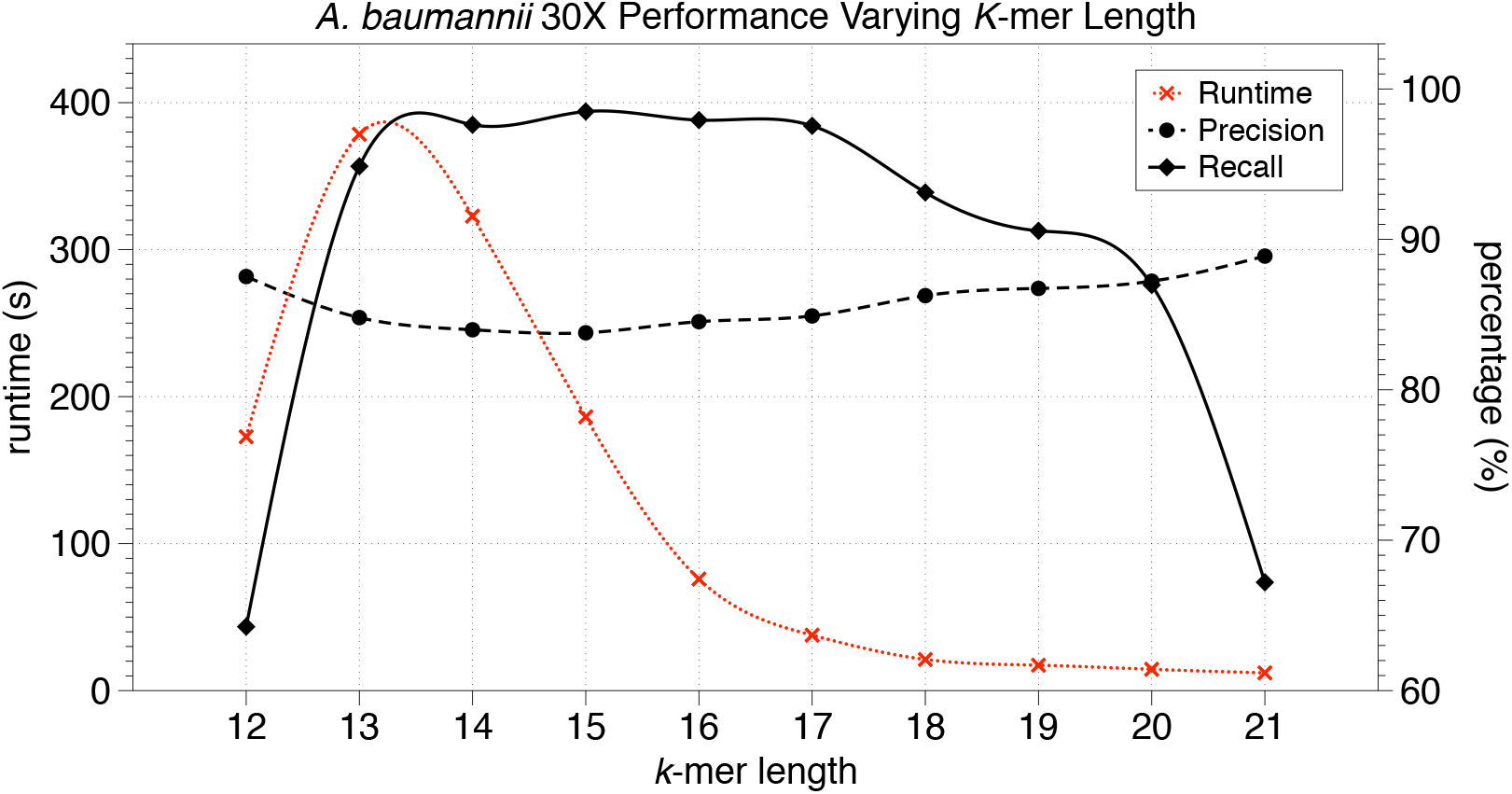
Recall, precision, and runtime varying the *k*-mer length for the *A. baumannii* 30X synthetic data set generated with an error rate of 15%.

### S3 Chernoff Bound for Overlap Detection

We will use a Chernoff bound (Chernoff *et al.*, 1952; Hoeffding, 1963) to define the value of *δ*. This way, we will prove that there is a very small probability of missing a true overlap of sufficient length *L* when using the (1 *− δ*)*φ* cutoff.

Let *Z* be a sum of independent random variables {*Z*_*i*_}, with *ϵ*[*Z*] = *μ_z_*; we assume for simplicity that *Z*_*i*_ ∈ {0, 1}, for all *i ≤ L*. The Chernoff bound defines an upper bound of the probability of *Z* to deviate of a certain quantity *δ* from its expected value. Specifically we use a corollary of the multiplicative Chernoff bound (Angluin and Valiant, 1979), which is defined for 0 *≤ δ ≤* 1 as:

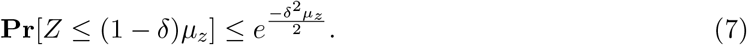

In order to obtain the Chernoff bound for the ratio *φ*, we consider a random variable *X*_*i*_ ∈ {−β, α} such that:

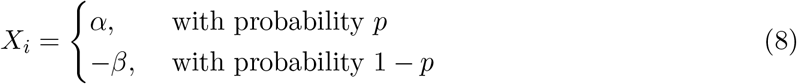

where *α, β >* 0 still are the values associated to a match and to a mismatch or a gap/indel in the scoring matrix, respectively; its expected value *E*[*X*_*i*_] is exactly equal to *φ* = *αp − β*(1 *− p*). Since the Chernoff bound is defined for a sum of independent random variables *Z*_*i*_ ∈ {0, 1}, we need to move from *X*_*i*_ ∈ {−β, α} to *Z*_*i*_ ∈ {0, 1}. Therefore, we define a new random variable *Y*_*i*_ = *X*_*i*_ + *β* as a linear transformation of *X*_*i*_, which can assume values *{*0*, α* + *β}*. Given *E*[*Y*_*i*_] = *E*[*X*_*i*_] + *β* = (*α* + *β*)*p*, we can normalize *Y*_*i*_ to obtain the desired random variable *Z*_*i*_:

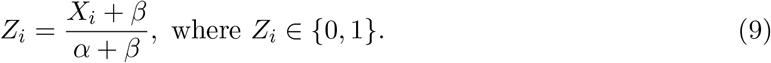

From the linearity of expectation, we have

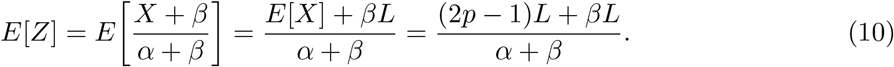

Substituting Eq. 9 and Eq. 10 in Eq. 7 and simplifying using our scoring matrix of *α, β* = 1, we obtain the final expression:

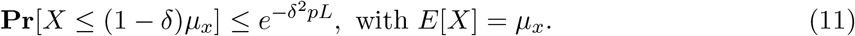

To interpret this bound, consider an error rate of 15%. Given two sequences that indeed overlap by *L* = 2000, the probability of their alignment score being below the mean by more than 10% (*δ* = 0.1) is *≤* 5.30 *×* 10^*−*7^.

### S4 Reliable Bounds Computation

Algorithm 2 and Algorithm 3 illustrates the pseudo-code used to compute the lower and upper bound, respectively, for our reliable range. The lower bound *l* is the smaller *m* value after which the cumulative sum exceeds a user-defined threshold *epsilon*, while the upper bound *u* is the largest value of *m* after which the cumulative sum exceeds *epsilon*.

### S5 Error Rate of the Input Read Set

The error probability *P*_*err*_ of a quality score *Q* is computed for each position of a read as proposed by Ono et al. (Ono *et al.*, 2012):

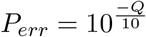

The error rate *e* of a given data set is then computed averaging the error probability *P*_*err*_ over the entire read set.

#### Algorithm 2

Reliable *k*-mer range: Selection of the lower bound (*l*)

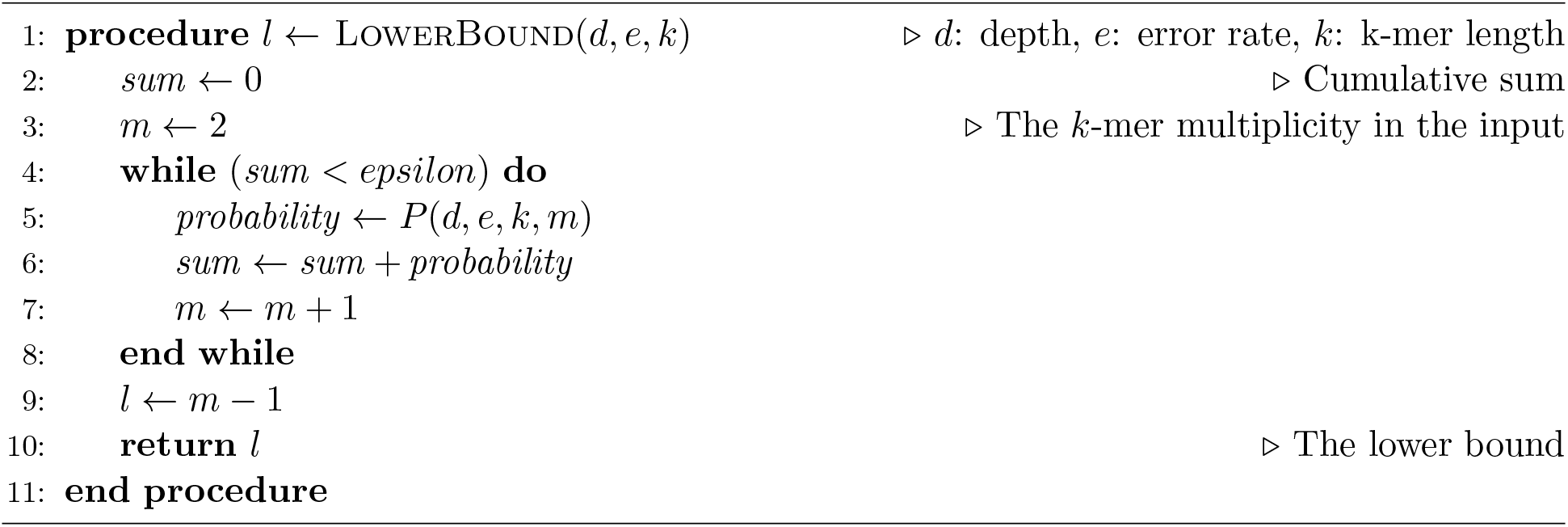

#### Algorithm 3

Reliable *k*-mer range: Selection of the upper bound (*u*)

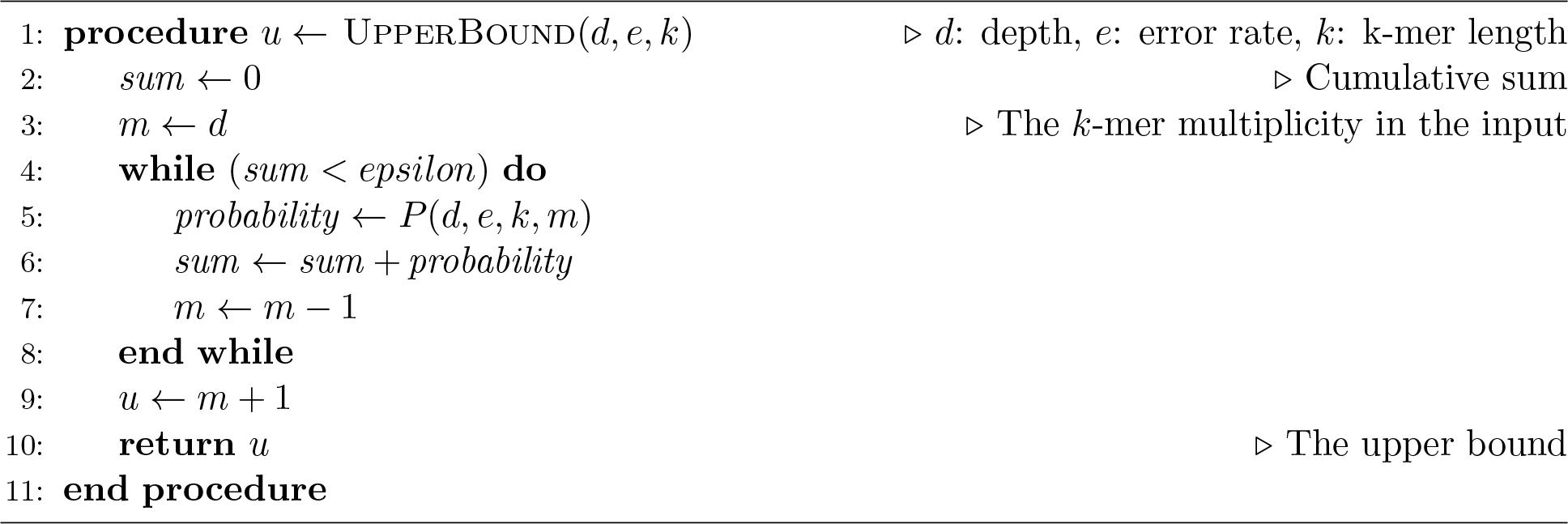

### S6 *X*-Drop Value

The choice of *k*, the x-drop value, and the alignment score threshold, all influence the final recall and precision. Our theoretical model for tuning these parameters assumes the typical read error-rates of high-throughput PacBio sequencing (10 *−* 15%). These errors are randomly distributed, and the majority of them are indels, erroneous insertions or deletions of nucleotides (Giordano *et al.*, 2017). Small values of *x* decrease the runtime but can potentially miss true overlaps in erroneous PacBio sequences. For example with *x* = 3, just 3 consecutive insertions on one sequence would cause the exclusion of a true overlapping pair from the output. Up to a certain point, increasing the value of *x* increases the number of true positives and make it easier to differentiate true alignments from false positives. Our SIMD seed-and-extend alignment with the adaptive band allows us to use *x* values *∈* [15, 100] without significant differences in the alignment runtime. However, higher values of *x* is not always better (Frith *et al.*, 2010). Consider two sequences of the type S1 = R1 *−* A *−* R2 and S2 = R2 *−* B *−* R2 where `R1` and `R2` are two repetitive regions of the genome. A large *x* value would allow the pairwise alignment algorithm to jump over `A` and `B` regions that are from different sections of the genome, potentially causing false positives depending on the relative length of `A/B` region to R1*/*R2 regions. For all our experiments, we used *x* = 50 with the exception of *C. elegans* 40X that we ran with *x* = 20 due to its lower error rate (*≈* 13%).

### S7 Output Format

BELLA outputs alignments in a format similar to BLASR/MHAP (Chaisson and Tesler, 2012; Berlin *et al.*, 2015), which outputs `.m4` files. For each pair of reads that pass both the overlapping and alignment stage filters, the output includes a line with the respective (a) the reads identifiers, (b) the number of shared *k*-mers, (c) the alignment score, (d) the overlap length estimate, (e) the strand information (*n* if the reads belong to the same strand, *c* if they are not, following DALIGNER (Myers, 2014) convention), (f) the start and end positions of the alignment in the first read, (g) the start and end positions of the alignment in the second read, and (h) the lengths of the reads.

### S8 Evaluation Procedure

We applied two different procedures to obtain values of recall and precision, one for real data and one for synthetic data. For each genome, a file containing the alignment positions of the input reads in the corresponding reference genome represents the *ground truth*.

The definition of ground truth is valid for both real and synthetic data. However, the ground truth generation procedure differs for the two categories of data as described in the following sections.

#### S8.1 Synthetic Data Set

Generating ground truth is simpler when dealing with synthetic data, and we accomplished it by running the read simulator of Pacific Biosciences, PBSIM (Ono *et al.*, 2012). The output of the simulator comprises the following files: one (or more, depending on the size of the genome) files in `fastq` format (i.e. the input file of BELLA and other software) and one (or more) files in the multiple-alignment format (MAF). The MAF file stores the information about where reads in the genome were synthesized; it is the equivalent of the single-mapped read-to-reference alignment ground truth generated for the real data sets using Minimap2. In this case, the exact location where a read was created is known, and therefore only single-mapped ground truth exists. We processed the MAF file (for synthetic data) and the two ground truth files (for real data) to extract the information required to compute the number of true overlaps: the read identifiers and the start and end positions in the genome.

The information stored in the ground truth was used to (a) compute the number of true overlaps, which constitutes the denominator in the formula to calculate recall and (b) store the reads information about their sub-reference sequence, and the start and end position in the reference. If the genome has more than one chromosome, we define the sub-reference sequence as the chromosome to which the read belongs; otherwise, the sub-reference sequence is the same for all the reads in a set. For each tool, we discarded self-paired reads (i.e., alignments of reads to themselves). Subsequently, we used the information stored in the ground truth to distinguish true positives from other read pairs. As previously defined, a read pair is considered a true positive if the two sequences align for at least *τ* = 2 kb, considering the start and end positions of their alignment in the reference genome (reported in the ground truth).

First, we estimated the overlap length from the information contained in the output of the considered tool, as described in Section S1. For this estimate, we used the length of the sequences, and the start and end overlap position on each sequence. If the pair does not satisfy this condition, but the pair is overlapping according to the ground truth, the pair it then counted as a false positive. If the pair does not appear in the ground truth, it is then counted as a true negative. If the overlap estimate is greater or equal to *τ*, then the pair is counted as a true positive if it is an overlap according to the ground truth, otherwise it is counted as a false positive. Given that a tool can report multiple locations for a given read-read pair, the overlap estimation is computed on the longest overlap a read-read pair presents in the output file of the considered tool. This choice maximizes the retention of true positives. This way, each read-read pair can contribute at most one to the total number of true positives, and one to the total number of identified overlaps. Diagram in Figure 6 summarizes the above described evaluation procedure.

**Figure 6:**
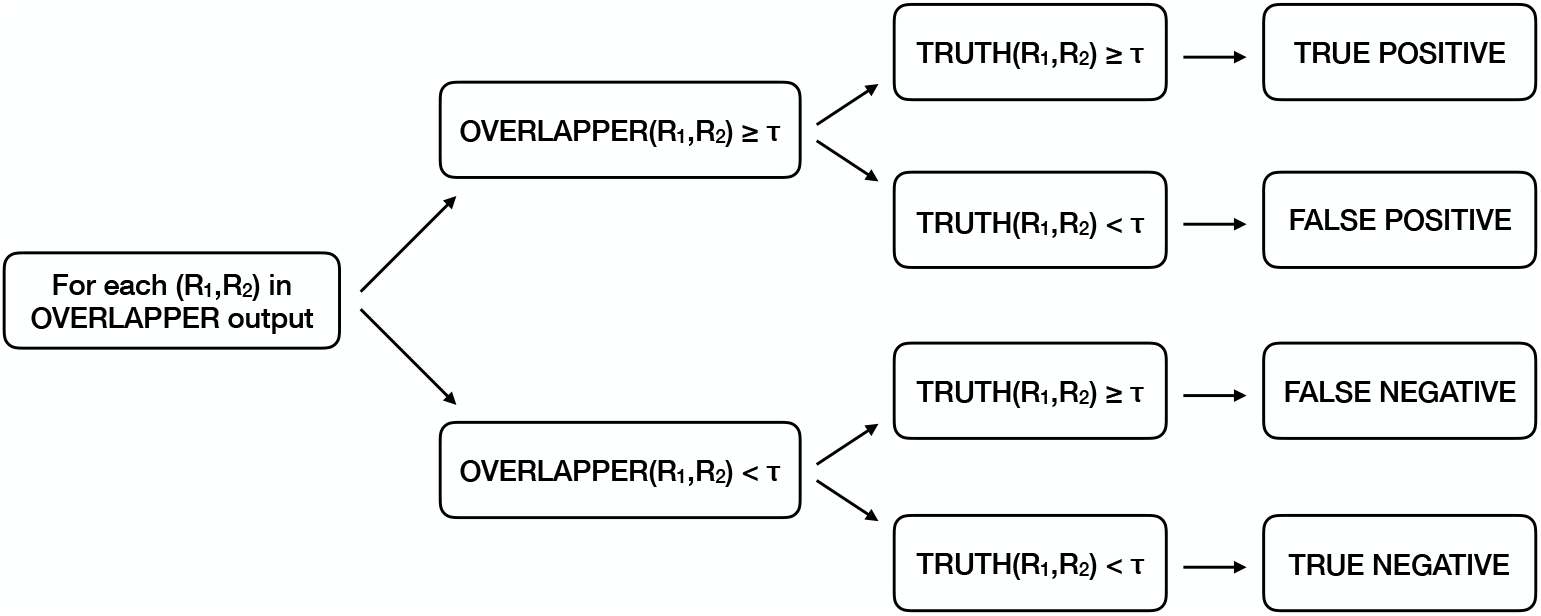
Diagram summarizing the evaluation procedure. (*R*_1_*, R*_2_) represents a pair of sequences, while *τ* is the threshold defining a true-positive overlap. *τ* is set to 2,000 bp in our default setting.

#### S8.2 Real Data Set

The procedure to generate the ground truth for real data is inspired by the one presented by Heng Li (Li, 2016), with one main difference. Our procedure generates two different ground truth files for real data (instead of one as in Li’s approach): a *single-mapped* file and a *multi-mapped* one; Li’s ground truth coincides with our single-mapped file. We used single-mapped ground truths to obtain the results shown in Table 2 and Table 3 in our main paper.

Following, we also consider the multi-mapped case because it is not always possible to be certain with real data, whether a read originated from one part of the genome or another, especially in the presence of repeats and high error rates. Hence, choosing a single mapping of a read could lead to the erroneous exclusion of valid read-to-reference alignments from the ground truth. It is our belief that a good overlapper/aligner should also be able to find correct overlaps that correspond to multi-mapped locations in the genome, as they could be useful for dealing with repeats in later stages of genome assembly. Therefore, the accuracy of each software was evaluated using the multi-mapped ground truth as well, as shown in Table 5. Naturally, the recall of any software would be lower (and its precision higher) if it were tested against multi-mapped ground truth, compared to if it were tested against single-mapped ground truth, since the former is a super-set of the latter.

**Table 5:** Recall, precision, and F1 score comparison (real data) with multi-mapped ground truth. The last column indicates whether the considered aligner does actual alignment or just overlap detection. Precision, recall, and F1 score are reported in percentage. Bold font indicates best performance and underlined font indicates second-best performance. BLASR result for *C. elegans* 40X is not reported as BLASR v5.1 does not accept `fastq` larger than 4 GB.

| Data Set | Overlapper | Recall | Precision | F1 Score | Time (s) | Alignment |
| --- | --- | --- | --- | --- | --- | --- |
| *E. coli* (Sample) | BELLA | 73.15 | **97.03** | <u>83.41</u> | 60.94 | Y |
|  | DALIGNER | <u>86.01</u> | 76.59 | 81.03 | **8.70** | Y |
|  | BLASR | 72.60 | 95.29 | 82.41 | 160.05 | Y |
|  | MECAT | 72.46 | 93.06 | 81.48 | 24.45 | N |
|  | Minimap2 | **87.44** | <u>93.47</u> | **90.35** | <u>16.57</u> | N |
|  | MHAP | 74.33 | <u>79.86</u> | 76.99 | <u>43.67</u> | N |
| *E. coli* | BELLA | 51.51 | 94.57 | 66.69 | 374.37 | Y |
|  | DALIGNER | 70.36 | 78.30 | <u>74.12</u> | **58.50** | Y |
|  | BLASR | 28.35 | **96.74** | 43.85 | 715.19 | Y |
|  | MECAT | 43.06 | <u>96.17</u> | 59.48 | <u>84.21</u> | N |
|  | Minimap2 | **70.02** | 90.72 | **79.03** | 107.76 | N |
|  | MHAP | 56.74 | 62.59 | 59.52 | 287.66 | N |
| *C. elegans* 40X | BELLA | 66.54 | <u>86.39</u> | <u>75.17</u> | 9,042.36 | Y |
|  | DALIGNER | 59.66 | 73.93 | 66.03 | 22,797.50 | Y |
|  | MECAT | 66.28 | **90.61** | **76.56** | **733.79** | N |
|  | Minimap2 | **88.00** | 42.25 | 57.09 | <u>1,733.26</u> | N |
|  | MHAP | <u>81.72</u> | 6.65 | 12.29 | <u>9,102.23</u> | N |

We mapped sequences against the reference genome by using both BWA-MEM and Minimap2 as we wanted to use the tool that minimizes the amount of incorrect (or missing) mapping positions; erroneous mappings would negatively affect our evaluation on real data sets. Table 6 suggests that Minimap2 may be a more suitable tool to map long reads to reference. Consequently, we evaluated the output of the overlappers against ground truth generated with Minimap2 to obtain the results.

**Table 6:** Comparison between BWA-MEM and Minimap2 in mapping against reference. The table shows recall and precision of whole set of tools when their outputs are compared against ground truth generated using Minimap2 or using BWA-MEM. The last two columns reports the percentage change of recall and precision when Minimap2 is used as ground truth instead of BWA-MEM. Brackets refer to negative values. BLASR result for *C. elegans* 40X is not reported as BLASR v5.1 does not accept `fastq` larger than 4 GB.

| Data Set | Tool | Minimap2 |  | BWA-MEM |  | Percent Variance |  |
| --- | --- | --- | --- | --- | --- | --- | --- |
|  |  | Recall | Precision | Recall | Precision | Recall | Precision |
| *E. coli* (Sample) | BELLA | 82.66 | 85.69 | 85.44 | 79.76 | (3.25) | 7.43 |
|  | DALIGNER | 89.97 | 62.62 | 90.48 | 56.70 | (0.56) | 10.44 |
|  | BLASR | 77.64 | 79.64 | 78.94 | 72.90 | (1.67) | 8.46 |
|  | MECAT | 78.41 | 78.71 | 80.30 | 72.58 | (2.41) | 7.79 |
|  | Minimap2 | 91.40 | 76.36 | 91.97 | 69.18 | (0.62) | 9.40 |
|  | MHAP | 79.71 | 66.93 | 80.93 | 61.18 | (1.53) | 8.59 |
| *E. coli* | BELLA | 65.08 | 71.22 | 69.06 | 63.25 | (5.76) | 12.60 |
|  | DALIGNER | 82.18 | 54.50 | 84.36 | 46.84 | (2.58) | 16.35 |
|  | BLASR | 35.41 | 72.01 | 37.69 | 64.16 | (6.44) | 10.90 |
|  | MECAT | 54.61 | 72.69 | 58.56 | 65.25 | (7.23) | 10.24 |
|  | Minimap2 | 80.68 | 62.30 | 82.03 | 53.02 | (1.67) | 14.90 |
|  | MHAP | 67.84 | 44.60 | 70.61 | 38.86 | (4.08) | 12.87 |
| *C. elegans* 40X | BELLA | 75.43 | 73.81 | 75.48 | 64.36 | (0.07) | 14.68 |
|  | DALIGNER | 62.81 | 58.66 | 60.27 | 49.04 | 4.04 | 19.62 |
|  | MECAT | 73.05 | 75.27 | 71.20 | 63.92 | 2.53 | 15.08 |
|  | Minimap2 | 94.13 | 34.06 | 91.49 | 28.84 | 2.80 | 15.33 |
|  | MHAP | 86.63 | 5.31 | 87.98 | 4.70 | (1.53) | 12.97 |

#### S8.3 Ground Truth Generation for Real Data

As with the original procedure presented by (Li, 2016), reads were aligned against the reference genome using BWA-MEM 0.7.17-r1188 (Li, 2013) with the following commands:

~~~
(1) bwa index <reference>.fasta
(2) bwa mem -x pacbio <reference>.fasta <input>.fastq >
<output>.sam
~~~

When using Minimap2 to map sequences to reference, the command is the following:

~~~
(3) minimap2 -ax map-pb <reference>.fasta
<input>.fastq > <output>.sam
~~~

The outputs of both BWA-MEM and Minimap2 were filtered in order to exclude non-mapped reads and alignments with mapping-quality lower that 10. The filtering was done using Samtools (Li *et al.*, 2009) with the following command:

~~~
(4) samtools view -h -Sq 10 -F 4 <output>.sam >
<multi-mapped>.sam
~~~

The output `<multi-mapped>.sam` obtained from the above operation constitutes the multi-mapped ground truth. The single-mapped ground truth `<single-mapped>.sam` was obtained by applying a second filtering step, using the following command:

~~~
(5) samtools view -h <multi-mapped>.sam | grep -v -e
’XA:Z:’ -e
’SA:Z:’ | samtools view -S > <single-mapped>.sam
~~~

through which command sequences mapped to multiple locations of the genome are removed, outputting only single-mapped sequences.

#### S8.4 Minimap2 versus BWA-MEM Reference Alignment

Table 6 shows recall and precision of each tool when its performance is evaluated toward a ground truth generated using both BWA-MEM and Minimap2. Precision significantly increases when Minimap2 is used to generate the ground truth, while the loss in recall is negligible. Overall, the results suggest that Minimap2 is more effective in mapping sequences to reference, reducing the number of missing alignments with respect to BWA-MEM. The author of both Minimap2 and BWA-MEM himself stated^3^ that Minimap2 is better than BWA-MEM to align sequences to reference when long-read data is used.

#### S8.5 Minimap2 and Simulated Data Reference Alignment

Table 7 compares the ground truth overlap cardinalities for synthetic data when such ground truth is generated using the real data procedure explained above against the actual synthetic data ground truth. Table 7 shows that Minimap2 is prone to overestimate the number of true overlaps with the exception of the *V. vulnificus* 30X data set. For comparison, we ran other two mapping software, BLASR (Chaisson and Tesler, 2012) and ngmlr (Sedlazeck *et al.*, 2018). Table 7 suggests that BLASR and ngmlr also tend to overestimate the ground truth cardinality.

**Table 7:** Comparison of ground truths generated using Minimap2, BLASR, and nglmr versus the *true* ground truth for synthetic data. The comparison is based on the number of overlaps reported as “true overlaps”. PBSIM indicates the *true* ground truth. BLASR’s result is not reported for *C. elegans* 20X as it exceeded the runtime limit of 48h.

| Data Set | Oracle | Overlaps Cardinality | Difference |
| --- | --- | --- | --- |
| *P. aeruginosa* 30X | PBSIM | 831,984 | - |
|  | Minimap2 | 953,680 | +14.62% |
|  | BLASR | 953,852 | +14.64% |
|  | ngmlr | 950,114 | +14.20% |
| *V. vulnificus* 30X | PBSIM | 692,576 | - |
|  | Minimap2 | 642,996 | -07.15% |
|  | BLASR | 934,410 | +34.92% |
|  | ngmlr | 610,358 | -11.87% |
| *A. baumannii* 30X | PBSIM | 577,970 | - |
|  | Minimap2 | 653,624 | +13.09% |
|  | BLASR | 654,832 | +13.30% |
|  | ngmlr | 647,632 | +12.05% |
| *C. elegans* 20X | PBSIM | 5,945,580 | - |
|  | Minimap2 | 6,777,820 | +14.00% |
|  | ngmlr | 6,659,504 | +12.01% |

#### S8.6 Output Translation to PAF Format

Miniasm assembler takes as input files in PAF format^4^. Thus, we had to translate each overlapper output format into PAF with the exception of Minimap2 which already uses that format.

The PAF format is defined as follow:

1. Query sequence name
2. Query sequence length
3. Query start (0-based)
4. Query end (0-based)
5. Relative strand: “+” or “-”
6. Target sequence name
7. Target sequence length
8. Target start on original strand (0-based)
9. Target end on original strand (0-based)
10. Number of residue matches
11. Alignment block length
12. Mapping quality (0 *−* 255; 255 for missing)

BELLA’s output format contains all the necessary information with the exception of the mapping quality (12), which is set to 255, and the number of residue matches *m* (10), which is estimated using the number of shared *k*-mers within a read pair. More formally,

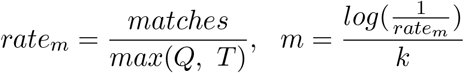

where *matches* indicates the number of shared *k*-mers within a read pair, *Q* and *T* correspond to the alignment length on the query and target sequence, respectively, and *k* is the *k*-mer length.

MECAT required the estimation of both (10) and (11). The alignment block length (11) defined as “the total number of sequence matches, mismatches, insertions and deletions in the alignment” was estimated using the start/end alignment information on the two sequences following the description in Section 3. MECAT’s output includes the percentage of identity between the two sequences, hence we used it to estimate the number of residue matches.

Finally, MHAP and BLASR use the `m4` format^5^. As for MECAT, the alignment block length (11) was estimated using the start/end alignment information on the two sequences. The `m4` format contains the alignment score that, however, cannot be directly used as an estimate for matches. Li^6^ writes that if the alignment is not performed, the alignment block length and the residual matches are still required, but their values may be highly inaccurate. Consequently, the number of residual matches was approximated by multiplying the alignment block length times the error rate.

### S9 Experimental Setting

The evaluation of competing software software was performed using MECAT 1.0, Minimap2 2.7, BLASR v5.1, MHAP 2.1.3, and DALIGNER 1.0, supported by DAZZDB v1.0. BLASR was run with the following settings: `--nproc 80 --maxLCPLength 16 --minMatch 12 -m 4 --bestn 50 --noSplit Subreads`, where `nproc` specifies the number of threads used to run BLASR, `maxLCPLength` indicates that the alignment of a read to the reference is stopped when its length reaches, 16 in our case (this command is useful when the query is part of the reference, which is the case when computing pairwise alignment for *de novo* assembly), `minMatch` is the minimum seed length, m indicates which file format of the output is used, `bestn` is the number of best alignments re-ported, and `noSplitSubreads` does not split input sequences at adapters; MHAP was run with: `--settings 3 --store-full-id --num-threads 80`, where 3 is the “sensitive mode” (Berlin *et al.*, 2015) and store-full-id stores sequences by their original identifier instead of a numerical index; MHAP was also run using 80 threads, as indicated by `num-threads`.

We collected the results on a dual-socket computer with two 20-core Intel Xeon Gold 6148 CPU (“Skylake”) processors, each running at 2.40 GHz with 384 GB DDR4 2400 MHz memory, using 2 threads per core (80 threads total). In order to run MHAP v2.1.3 on *C. elegans* 40X and *C. elegans* 20X, we increased the Java heap space to fit our largest machine, since the default 32 GB setting (used for all the other data sets) led to out-of-memory failures.

Finally, DALIGNER results are only reported for real PacBio data because it requires single cell PacBio sequencing data (Myers, 2014). For all of the simulated data sets, DALIGNER failed-fast, producing the associated error, `“Pacbio header line format error”`. Our attempts to reformat the simulated data (to “fool” DALIGNER) failed with the same error.

### S10 Memory Usage

Table 8 presents a comparison of the relative memory consumption of each software tool on two representative data sets. The memory usage information was collected with Valgrind (Nethercote *et al.*, 2006) on dedicated dual socket compute nodes, with up to 128GB RAM and a 16-core Haswell CPU per socket. The Valgrind version was `3.15.0`, and the following command was used: `valgrind --tool=massif --pages-as-heap=yes`. Each tool was compiled on the described platform with -g (as required by Valgrind); otherwise, no changes were made in the compilation procedure from that used for the results in Section S9, with one exception. MHAP was compiled with the addition of `java -Djava.compiler=NONE`. DALIGNER encountered early segmentation faults when run with 2 threads per core (`-T64`), due to its per-thread memory demands. The results shown were collected with `-T32`.

**Table 8:**
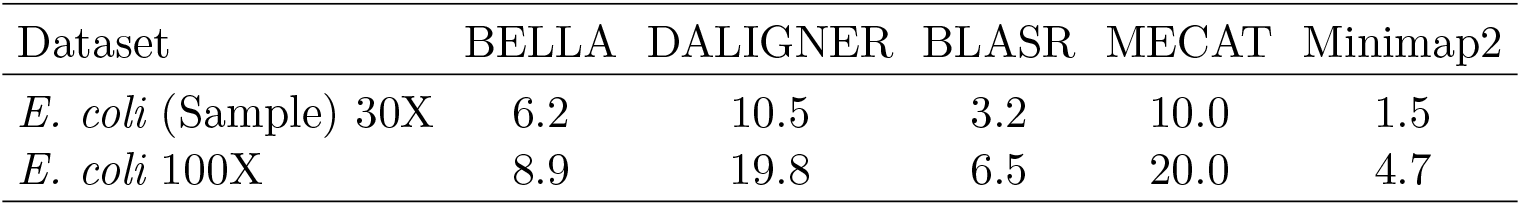
Peak memory usage for two representative data sets. The numbers are reported in GB. MHAP’s results are not reported as it exceeded the maximum time limit of 48h.

| Dataset | BELLA | DALIGNER | BLASR | MECAT | Minimap2 |
| --- | --- | --- | --- | --- | --- |
| *E. coli* (Sample) 30X | 6.2 | 10.5 | 3.2 | 10.0 | 1.5 |
| *E. coli* 100X | 8.9 | 19.8 | 6.5 | 20.0 | 4.7 |

### S11 Performance Breakdown of BELLA

Figure 7 illustrates the runtime breakdown of BELLA as a percentage of the total runtime. Pairwise alignment dominates our runtime, while sparse matrix construction, which includes the creation of both **A** and **A**^T^, and multiplication take only a tiny percentage of our computation, proving the efficiency of our approach for overlap detection.

**Figure 7:**
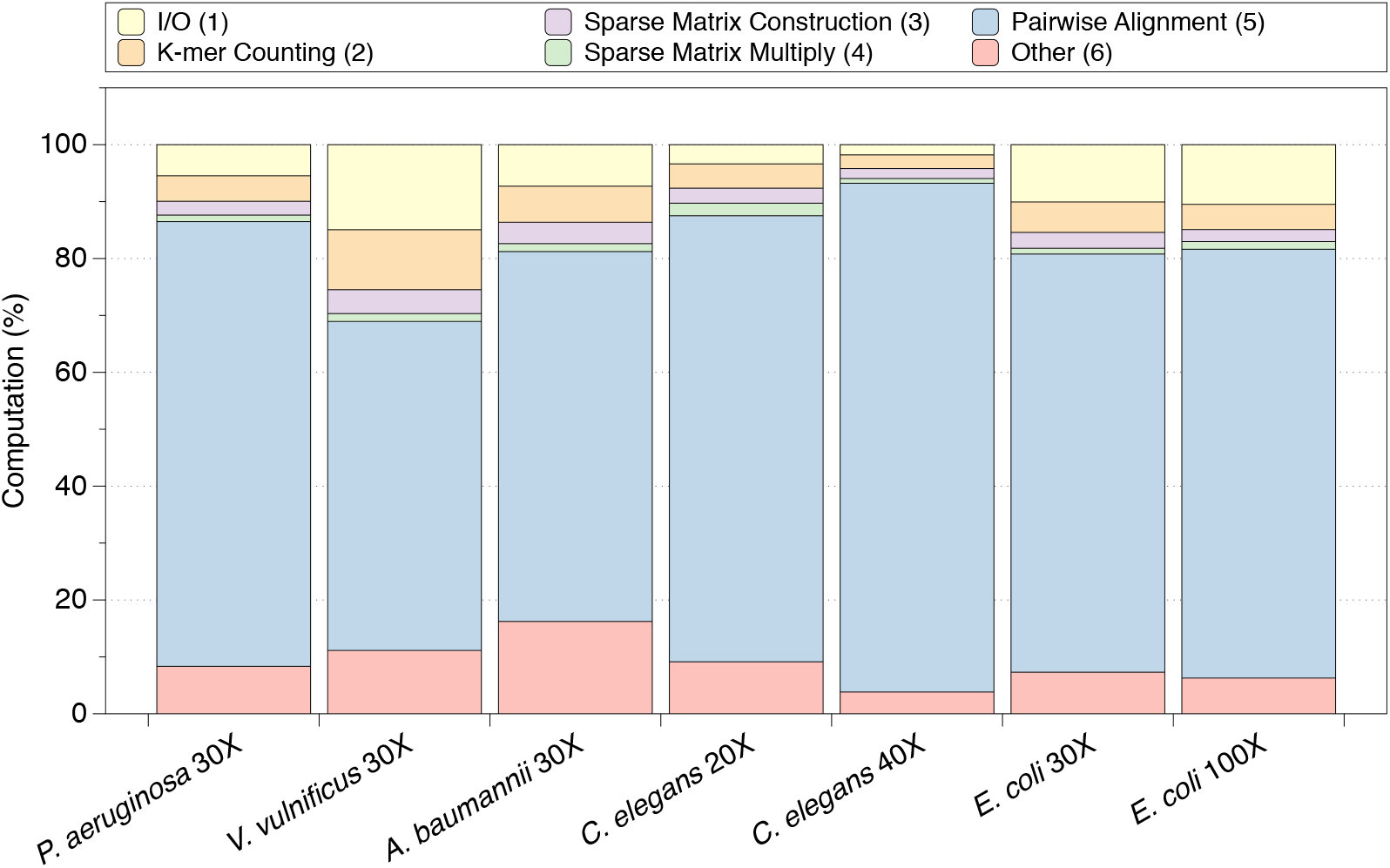
BELLA’s runtime breakdown. The runtime is divided into (from the top of the bar): (1) I/O, (2) *k*-mer counting using a Bloom filter and a multithreaded hash table, (3) sparse matrix construction, (4) sparse matrix multiplication, which is BELLA’s overlap detection, (5) nucleotide-level pairwise alignment, and (6) other.

Interestingly, sparse matrix multiplication and semiring abstraction could offer a path for efficient parallelization of many applications in computational biology other than overlap detection (Jain *et al.*, 2019).

### S12 Scalability

Figure 8 shows the strong scaling curves of BELLA for the representative *P. aeruginosa* 30X data set to measure its parallel performance. The data was collected on the same compute nodes as those described in Section S9. The 80-way thread parallelism is achieved by exploiting hyper-threading as each node has 40 cores and each core supports 2 hyper-threads.

**Figure 8:**
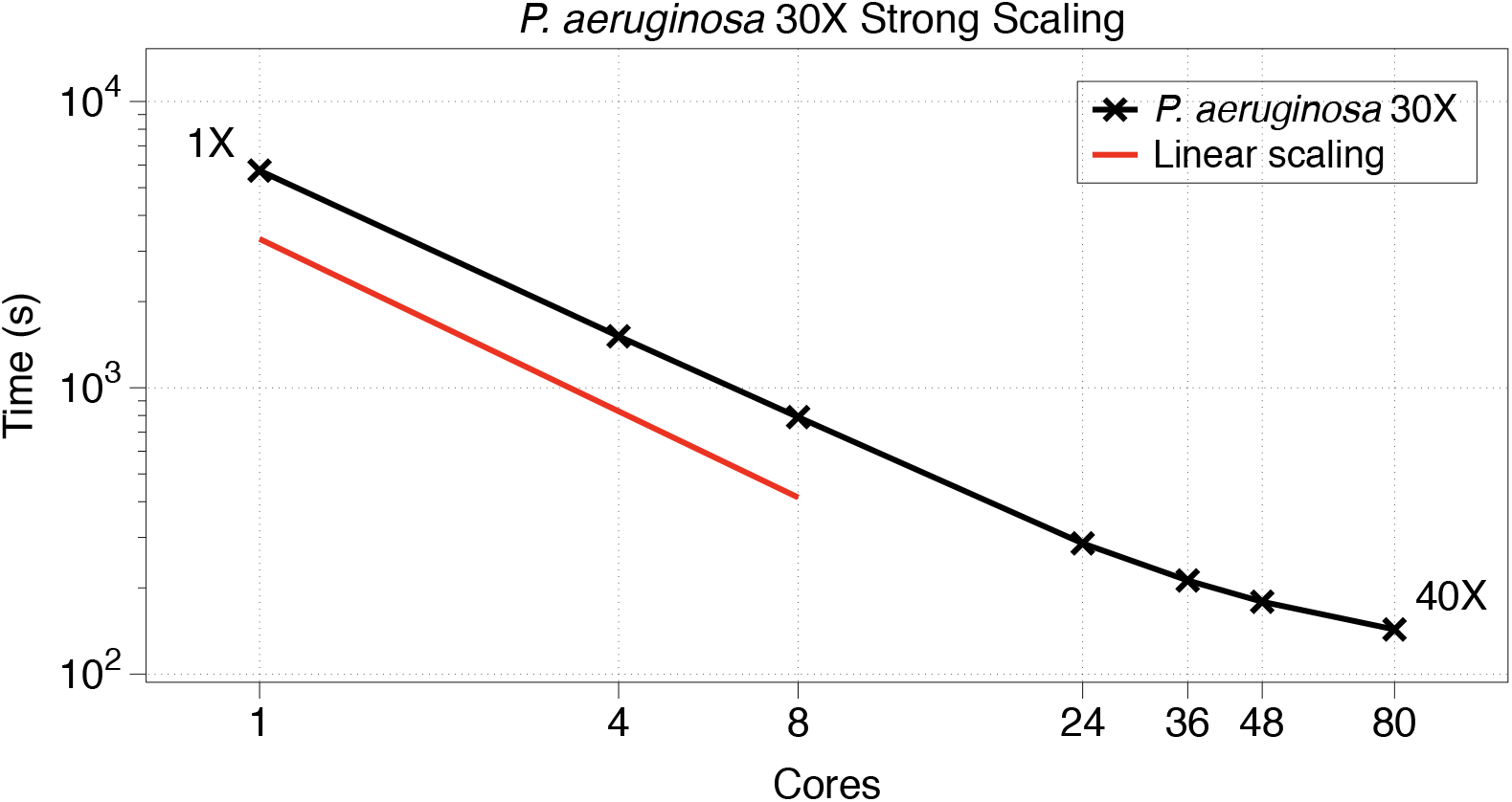
Scaling of BELLA with increasing number of cores on a dual-socket computer with two 20-core Intel Xeon Gold 6148 CPU (“Skylake”) processors, each running at 2.40GHz with 384GB DDR4 2400 MHz memory.

In strong scaling, the problem size stays fixed but the number of processing elements is increased. In strong scaling, a program is considered to scale linearly if the speedup is equal to the number of processing elements used, corresponding to the number of threads in our case. Given the amount of time to complete a given run using one thread is *t*_1_ and the amount of time to complete the same run with *n* threads is *t*_*n*_, we can define the strong scaling efficiency (as a percentage of linear) as:

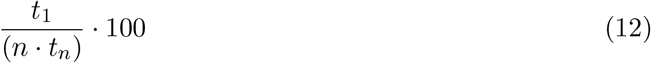

BELLA with 80-way thread parallelism had a strong scaling efficiency of 50.07% and a speedup of 40*×* for this representative data set.

## Footnotes

1 https://github.com/PacificBiosciences/pbbioconda/issues/46

2 Like other parameters of BELLA, this can be adjusted in the command line to detect shorter overlaps, potentially resulting in lower precision and higher recall

3 https://lh3.github.io/2018/04/02/minimap2-and-the-future-of-bwa

4 https://github.com/lh3/miniasm/blob/master/PAF.md

5 https://github.com/PacificBiosciences/blasr/wiki/Blasr-Output-Format

6 https://github.com/lh3/miniasm/blob/master/PAF.md

